# Time and mode of Culicidae evolutionary history

**DOI:** 10.1101/871145

**Authors:** Alexandre Freitas da Silva, Laís Ceschini Machado, Marcia Bicudo de Paula, Carla Júlia da Silva Pessoa Vieira, Roberta Vieira de Morais Bronzoni, Maria Alice Varjal de Melo Santos, Gabriel Luz Wallau

## Abstract

Mosquitoes are insects of medical importance due their role as vectors of different pathogens to humans and other mammals. There is a lack of information about the evolutionary history and phylogenetic positioning of the majority of mosquitoes species. Here we sequenced the mitogenomes of mosquitoes species through low-coverage sequencing and data mining. A total of 37 draft mitogenomes were assembled representing 11 genera and 16 of those were sequenced for the first time. The recovered mitogenomes showed a coverage breadth average of 81.24%. Most of the species were clustered in monophyletic clades with other members of their own genus with exception of the Aedini tribe which was paraphyletic corroborating other findings. We established for the first time the monophyletic status of eight species from the tribe Mansoniini including both *Coquillettidia* and *Mansonia* genus and established the basal positioning of Aedeomyiini and Uranotaeniini tribes regarding the Culicinae subfamily. Molecular clock dated the Culicidae family emergence around 273 MYA and the split between *Anophelinae* and *Culicinae* subfamily around 182 MYA in the Jurassic period. Low-coverage sequencing is effective to recover mitogenomes, establish phylogenetic knowledge generating basic fundamental information to the understanding of the role of these species as pathogen vectors.

## Introduction

Mosquitoes compose a larger group of insects from the Culicidae family. There are around 3,567 valid species classified into two subfamilies (*Anophelinae* and *Culicinae*) and 41 genera (http://mosquito-taxonomic-inventory.info/ accessed on 21 Oct., 2019). The large majority of mosquitoes species have an atropofilic behaviour towards reptiles and mammals including humans ^1^. Because of that they can transmit many pathogens such bacteria ^2^, malaria protozoa (OMS, 2018), filarial worms ^3^ and arboviruses ^4^. Mosquitoes are responsible for the transmission of pathogens that cause outbreaks and epidemics annually in the tropical region, but the current globalization process and land use change are rising human-mosquito contact allowing the emergence of new mosquito-borne disease ^5–7^. Several of the new emerging pathogens arose from forested environments where they circulate in a sylvatic cycle between wild animals and arthropod vectors species such as mosquitos ^8^. Although there are abundant evidence that spillover occurs from sylvatic to urban environments, we know very little about the sylvatic cycle of these pathogens including the vector species that transmit them in their natural environment ^9^. Therefore, basic knowledge about vector evolution and ecology is highly needed to better understand their role in the transmission cycle of pathogens ^5,10,11^.

The huge improvement in nucleic acid sequencing platforms in the last decade has allowed an explosion of genomic information from a wide range of species. Mitogenomes, the entire mitochondrial genome, have been widely used as a target molecule to elucidate different aspects of metazoa species evolution such as population dynamics and phylogenetic relationships^12^. Complete mitogenomes are reliable tools to be used as a molecular marker in ecological and evolutionary studies because they provide genes with different evolutionary rates such as the most conserved rRNA genes (12S and 16S), the intermediate ND1-6 genes and the fast evolving Cytochrome oxidase subunit I (COI) gene, the most used molecular marker for species identification, allowing an accurate establishment of both ancient and recent speciation events ^13–15^. In addition, mitogenomes have a uniparental heritage, high copy number per cells and single-copy genes which facilitates DNA recovery and phylogenetic analysis ^16–18^. Recently, some studies have sequenced a larger number of mitochondrial genomes from different mosquito species, but they are mostly focused on *Anopheles* species ^19–21^. Mosquito mitogenomes are structurally conserved following the metazoa gene number and order, with few exceptions, showing 37 genes comprising 13 protein coding genes, 22 tRNAs and 2 rRNA genes ^22–24^. Its size range varies from 14.820 bp for *An. maculatus* to 16.790 bp for *Ae. aegypti* (NCBI, https://www.ncbi.nlm.nih.gov/genome/browse#!/organelles/culicidae accessed on 21 oct., 2019).

Mitogenome sequencing has been a hard task using the first generation of sequence platforms based on the Sanger method. The first mosquito mitogenome have been obtained after laborious steps such as mitochondria purification followed by DNA extraction, cloning and Sanger sequencing of several fragments ^25,26^. Today there are a number of alternative approaches available to obtain mitogenomes which was only possible due to the improvement of second and third generation of sequencing platforms. Most of these strategies are based on PCR/Long Range PCR coupled with Next-generation sequencing (NGS), shotgun Whole Genome Sequencing or mitogenome sequencing through RNA-Seq data ^27,28^. Other approaches available allow the recovery of mitogenomes by PCR amplification from environment samples and pooled DNA and mitogenome recovery from low-coverage sequencing ^29–31^. Moreover, a number of bioinformatics tools were developed to specifically assembly and annotate mitogenomes ^32–36^

Most of the available mitogenomes belong to *Anopheles* species with much fewer genomes from *Culex, Aedes* and other genera such as *Haemagogus, Bironella, Sabethes,* and *Lutzia* ^19–21,24,37^, but there is no available molecular data for the large majority of the species. Aiming to contribute with this basic and fundamental knowledge we performed low-coverage sequencing and data mining on already published Culicidae SRA data in order to characterize the mitogenomes from different species and genera and better understand the evolutionary history of mosquitoes focusing in Culicinae subfamily. Overall, we reconstructed and positioned 37 mitogenomes, 35 of them for the first time, representing 11 genera and performed molecular dating of speciation time of all common ancestors.

## Material and methods

### Mosquito sampling and taxonomic identification

Mosquito samples were collected in remnants of the brazilian Atlantic forest and from the South border of the Amazonian forest Brazil. Three locations were sampled at Pernambuco state, Parque Estadual Dois Irmãos (8°00′43.3″S 34°56′40.7″W) at the Recife municipality; Reserva Ecológica de Carnijó (8°08′20.7″S 35°04′47.3″W) at Moreno municipality; and Aldeia at Camaragibe municipality. Additionally, one sampling was performed at Sinop municipality, Mato Grosso state. Different sampling methods were employed aiming to collect a large diversity of species. Diurnal sampling was performed with aspirators (Horst model) and entomological nets, larvae and pupae were collected on water pools and plant holes. Nocturnal sampling was performed using CDC-light traps and BG-Sentinel were used to sample mosquitoes attracted by light and odorants. The specimens were transported alive to the Entomology department of Aggeu Magalhães Institute - Oswaldo Cruz Foundation (IAM/FIOCRUZ), immature specimens were maintained in liquid water and feed with cat food (Friskies®) until the emergence of adults. Adults mosquitoes were separated per morphological groups, were dry stored in silica at room temperature until taxonomic identification. Taxonomic keys for neotropical Culicidae were used for species identification ^38,39^. Besides the collection performed in this work we included *Aedes taeniorhynchus* and *Aedes scapularis* samples ceded by collaborators of the Entomology department of IAM sampled respectively at São Luis municipality, Maranhão state and Juazeiro municipality, Bahia state. All collections were authorized by the regulatory organ - SISBIO under the license number: 58716-1.

### DNA extraction and sequencing

The specimens were macerated in ultrapure water using 40ul/individual in single or pooled samples (Supplementary table 1) according to the number of specimens collected per species. Both male and female individuals and samples from different collection points were included in the pools. Total DNA extraction were performed using ethanol precipitation method^40^ and QIAprep Spin Miniprep extraction (QIAGEN) in order to improve mitochondrial DNA by enrichment as suggested by Quispe-Tintaya et al. (2013). All samples were assessed by quality and purity with NanoDrop 2000 (Thermo Scientific) and quantified through Qubit® dsDNA HS (High Sensitivity Assay) kit. The DNA library was prepared using the Nextera XT library preparation kit following the recommendations of the manufacturer (Illumina, San Diego, CA, USA). DNA library was sequenced on the Illumina Miseq platform using a paired-end approach of 75 bases with Reagent Kit V3 of 150 cycles.

### Dataset construction

A search on National Center for Biotechnology Information (NCBI) was performed to recovery already characterized mitochondrial genomes (mitogenomes) from mosquitoes representing different genus (Supplementary table 2). Besides, we also searched on the SRA database for mosquitoes raw sequence reads (Whole genome sequencing and RNA-Seq) available up to November of 2018 representing species that had no mitogenome available at the time (Supplementary table 3).

### Quality control of sequences

The raw reads (sequenced in this study and recovered from SRA) were checked for quality using FastQC program (http://www.bioinformatics.babraham.ac.uk/projects/fastqc/) and results were summarized on MultiQC tool ^41^. Based on the excellent quality of our sequenced raw reads they were not trimmed (Supplementary figure 1) but, all SRA libraries were trimmed using the Trimmomatic tool v 0.35 ^42^ to remove adapters and ensure that the quality of sequences (Phred score >20).

### Mitogenome assembly and annotation

The mitogenomes were assembled using a baiting and iterative mapping approach implemented in MITObim 1.9 ^34^. Different mosquito mitogenome were used as reference genome for the first capturing of reads considering the closest mitogenome available to each species analysed. SRA reads were assembled using MITObim default parameters (*-kbait* parameter=31). While we used a combination of parameters to generate a consensus sequence for the sequenced species. A first assembly was performed with -*kbait* =15 followed by a second assembly step using -*kbait* = 31. The final consensus assembly was composed by the consensus of the two assemblies that was then checked with well characterized mitogenomes to correct any potential assembly errors (ex. the assembly of non alignable regions between mitogenomes). To assess the average coverage depth of each mitogenome, the reads were mapped against the assembled mitogenomes through the *MIRAbait* module from MIRA sequence assembler software^43^. Automatic gene annotation were performed on MITOS2 web server (available on http://mitos2.bioinf.uni-leipzig.de/index.py) ^35^ based on invertebrate genetic code against the metazoan Refseq 81. Comparative genomics maps were built using *Ae. aegypti* mitogenome (Accession number: NC_010241.1) as reference in BRIG (BLAST *Ring Image Generator*) ^44^.

### Evolutionary analysis

Nucleotide mitogenomes sequences characterized here, were aligned by MAFFT v 7.0 tool ^45^ with already characterized mitogenomes recovered from databases (Supplementary table 2). The non aligned sites were removed using GBLOCKS tool v. 0.91b - default parameters, with exception for the allowed gap positions that was set as with the “half” option ^46^ and final alignment was visualized on Aliview ^47^. The evolutionary analysis were performed based on four possible alignment approaches: I) Complete nucleotide mitogenome alignment sequences, II) Partitioned nucleotide sequence of protein coding genes, III) Partitioned predicted amino acid sequences from coding regions and IV) Concatenated alignment of amino acid sequences. Nucleotide substitution saturation analysis was performed for each nucleotide gene alignment in DAMBE software ^48^ evaluating 1st + 2nd and 3rd codon position separately through the Xia et al. test ^49^. Nucleotide substitution models for I and II alignments were obtained with Smart model selection (SMS) implemented on PhyML webserver ^50^. Protein evolutionary models were assessed for III and IV alignments using Prottest 3.4.2 ^51^. All phylogenetic analysis were based on a Bayesian Markov Monte Carlo approach (MCMC) performed on BEAST 1.8.4 package ^52^ in order to infer the topology of Culicidae family and the speciation time of the common ancestor of clades in million years. In order to obtain calibration points for the molecular clock analysis, we performed a literature search to obtain fossil dates representing the different Culicidae clades. Although there are several potential calibration points to the Culicidae tree we only kept the ones supported by fossil evidence. We used four calibration points representing the Diptera order, Culicidae family, Anophelinae and Culicinae subfamily (Supplementary table 4).

Bayesian analysis was performed with at least three independent runs of 150 million generations sampling at each 1000 trees, for each alignment dataset. The lineage through time analysis and Effective sample size (ESS) were evaluated by Tracer 1.7.1 ^53^ and reached 200 for most of the important parameters for dating and tree likelihood. The analysis were performed under an uncorrelated relaxed molecular clock using a lognormal distribution and a Birth-Death model process of speciation as Tree Prior. For the complete mitochondrial genomic alignment (alignment I) the GTR+G+I evolutionary model was used. For the partitioned gene analysis (alignment II) and partitioned predicted amino acids (alignment III) each partition was set with a specific evolutionary model as previously described (Supplementary table 5). Besides the partitioned gene analysis we also performed a more robust analysis based on the nucleotide saturation of each gene taking into account the partition of codons positions where the 1st and 2nd codon positions were split from the 3rd codon position. The concatenated protein analysis was performed under the mtREV+G+I evolutionary model. The posterior probability tree for each alignment dataset were built combining the three independent runs of each analysis with LogCombiner program applying a burn-in of 25% and the consensus credible tree was obtained through the TreeAnnotator program. The timescale trees were plotted with Phyloch package version 1.5-3 (available on http://www.christophheibl.de/Rpackages.html) from R programming language. Tres topologies comparison were performed by plotting tanglegrams using the Dendextend R package ^54^.

## Results

### Sequencing and mitogenome characterization

The sequenced mosquito samples generated a total of 84.2 million paired-end reads representing the seventeen species and eight genera (*Aedeomyia, Aedes, Coquillettidia, Culex, Mansonia, Psorophora, Trichoprosopon* and *Uranotaenia*). The amount of generated reads ranged from 1.1 million reads for *Uranotaenia pulcherrima* to 11.3 million reads for *Aedes taeniorhynchus* (Table 1). Searching on the SRA database we include other twenty mosquito species raw read datasets for mitogenome characterization representing six genera (*Aedes, Anopheles, Culex, Psorophora, Tripteroides* and *Toxorhynchites*). We characterized 35 new draft mitogenomes for Culicidae family in total. Sixteen out of seventeen mitogenomes were sequenced for the first time considering only the field-sampled species (*Ad. squamipennis, Ae. scapularis, Ae. taeniorhynchus, Cq. albicosta, Cq. chrysonotum, Cq. hermanoi, Cq. juxtamansonia, Cq. venezuelensis, Cx. amazonensis, Cx. corniger, Ma. wilsoni, Ma.humeralis, Ma.titillans, Ps. cingulata, Tr. digitatum, Ur. pulcherrima)*. In summary, the newly characterized mitogenomes represent eight Culicidae genera (*Aedeomyia, Coquillettidia, Mansonia, Psorophora, Trichoprosopon, Tripteroides, Toxorhynchites,* and *Uranotaenia*) that had no mitochondrial genome data available at the moment.

**Table 1.**
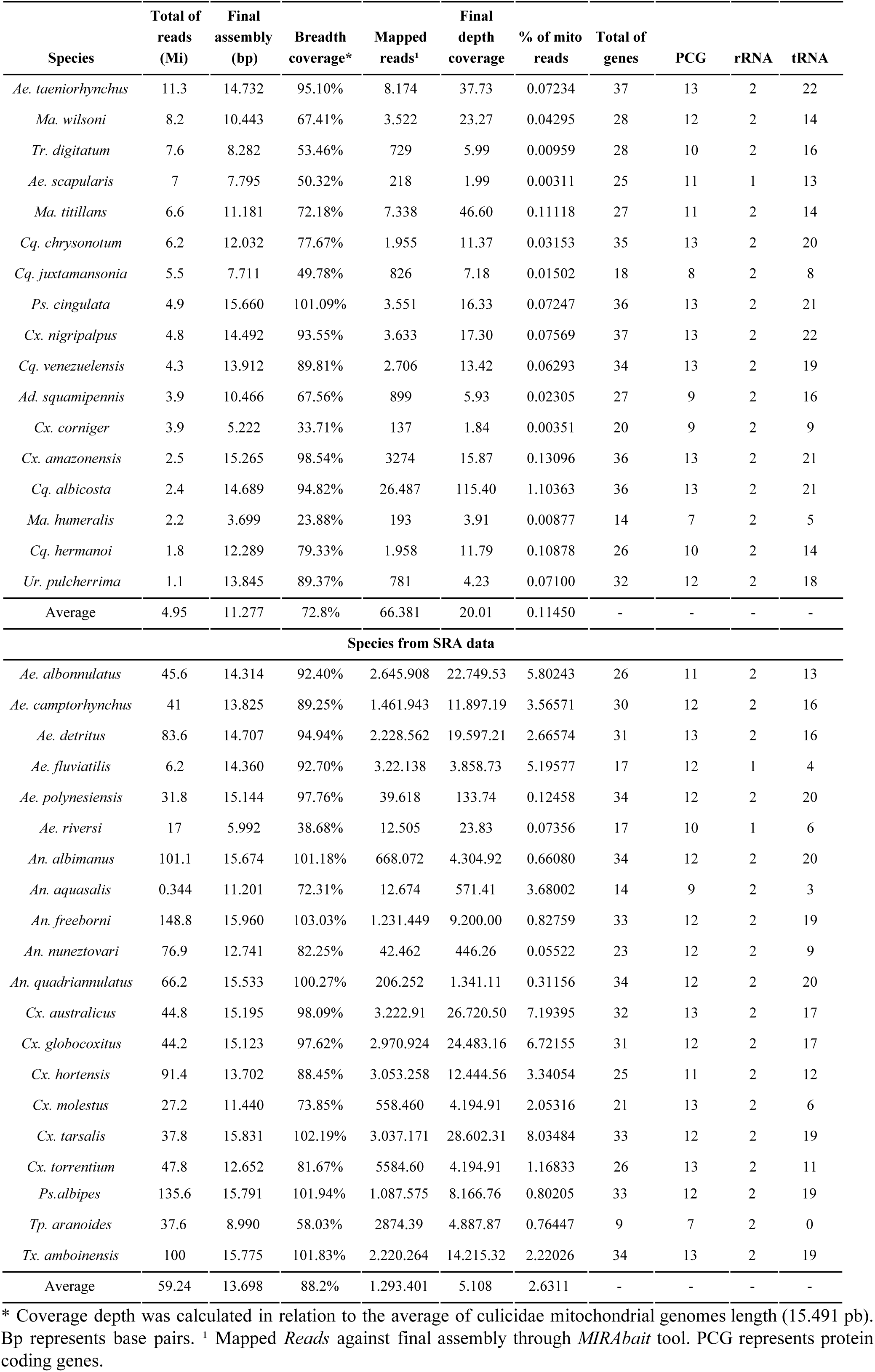
General statistics for sequenced draft mitogenome.

The coverage breadth of the sequenced draft mitogenomes ranged from 3.699 bp to 15.660 bp for *Ma. humeralis* and *Ps. cingulata* respectively (Table 1, Figure 1) with and average of 72.80% and a coverage depth average of 20.01 fold (Table 1). Annotation of the protein coding genes (PCG) identified in the field-collected mosquitoes ranged from 7 to 13. All seventeen mitogenomes showed the two rRNA genes except *Ae. scapularis* genome. While tRNAs annotation ranged from 5 to 21 genes, except for *Ae. taeniorhynchus* and *Cx. nigripalpus* that showed all tRNAs genes (Table 2). Although some PCGs were not assembled we could annotate the barcode Cytochrome oxidase I gene in all seventeen mitogenomes (Supplementary table 6). The mitogenomes characterized from SRA data showed a coverage breadth ranging from 5.992 to 15.960 bp for *Ae. riversi* and *An. freeborni* respectively (Figure 2). In general those assemblies showed an average coverage breadth of 88.42% and from 9 to 34 out of 37 mitochondrial genes were annotated with MITOS (Table 1). Although some of the SRA data came from RNA-Seq we were able to identify almost all PCGs of these mosquito species. PCGs annotation ranged from 7 for *Tp. aranoides* to 13 for some species (Supplementary table 6, Figure 2).

**Figure 1.**
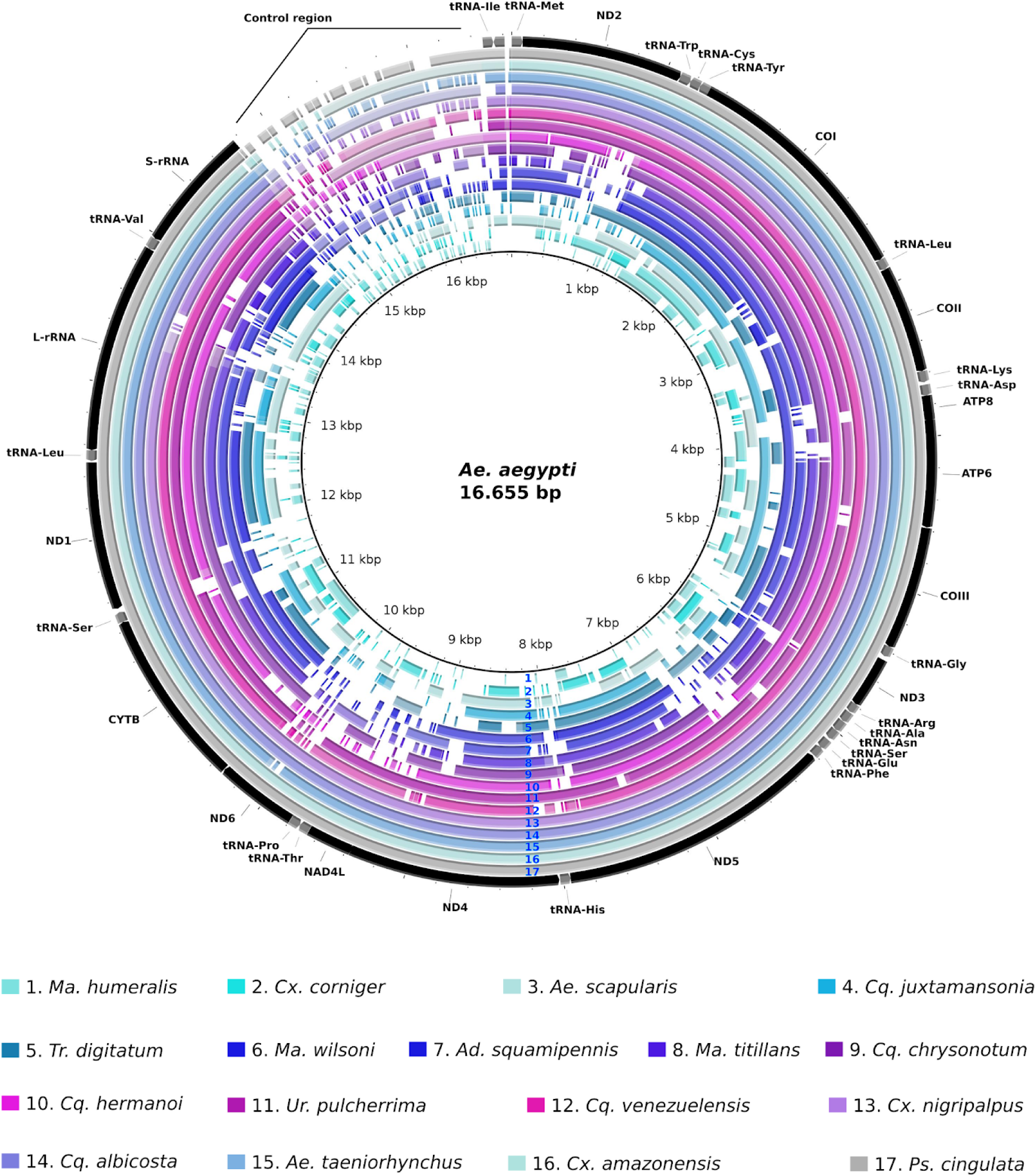
Comparative map of mitogenomes sequenced in relation to *Ae. aegypti* mitochondrial genome (NC_010241.1).

**Figure 2.**
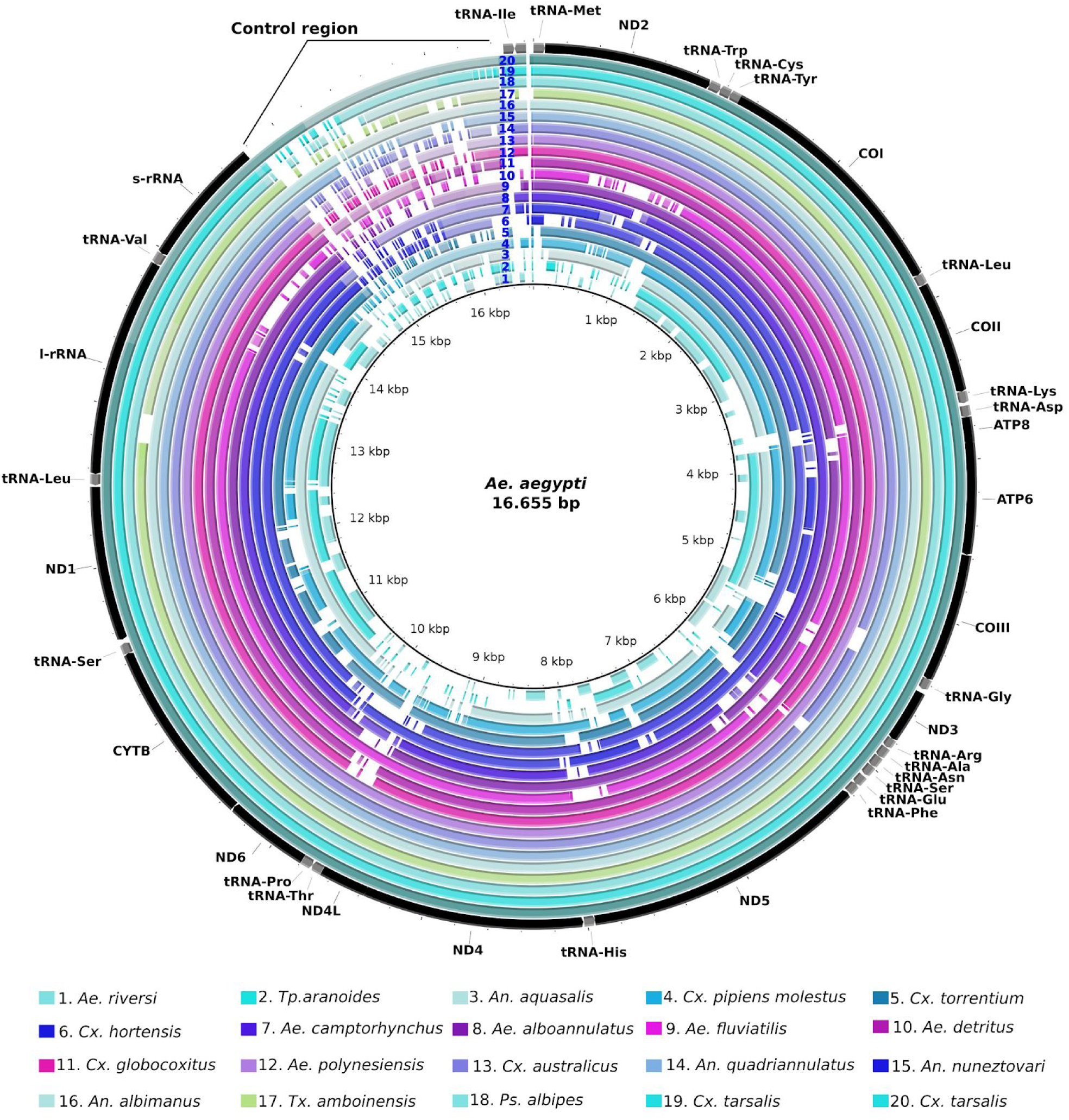
Comparative map of mitogenomes characterized from SRA data in relation to *Ae. aegypti* mitochondrial genome (NC_010241.1).

### Evolutionary analysis

In order to establish the phylogenetic relationship of Culicidae family we performed alignments with nucleotide and amino acid datasets taking into consideration or not gene/protein partitioning. Since several genes showed nucleotide saturation at the third codon position we also partitioned the codons of each PCG (Supplementary file 1). Topology of the phylogenetic trees built with those different alignments were mostly in agreement but also showed some clade grouping differences. *Aedeomyia, Uranotaenia* and *Toxorhynchites* genus were basal to the Culicinae subfamily but different grouping between them and uncertainty in the deep branching pattern appeared between analysis (Supplementary figure 2 and 3). Moreover, a number of intra genus incongruences between trees was also observed in the *Culex, Anopheles* and *Aedes* genera (Supplementary figure 2 and 3). The comparison between amino acid phylogenies against partitioned codon PCG phylogeny showed a similar topology supporting the basal position of some clades as *Uranotaenia* and *Aedeomyia* from Culicinae subfamily. Nevertheless, some branches within genera such as *Coquillettidia*, *Culex, Aedes* and *Anopheles* showed different positions (Supplementary figure 3).

The Culicidae family split from other dipterans during Permian period around 273 million years ago (MYA) (Figure 3, node A and Supplementary table 7). While the most recent common ancestor of the Culicidae family emerged in the Jurassic period around 182 MYA with the *Anophelinae* and *Culicinae* subfamilies origin (Figure 3, node B). In the subfamily *Anophelinae,* the *Chagasia* genus showed basal to *Bironella* and *Anopheles* genera with speciation in the Cretaceous period around 145 MYA (Figure 3, node C). The last two genera showed speciation times from 110 MYA until the last 2 MYA in the *gambiae* species complex (Figure 3, nodes D and E, respectively).

**Figure 3.**
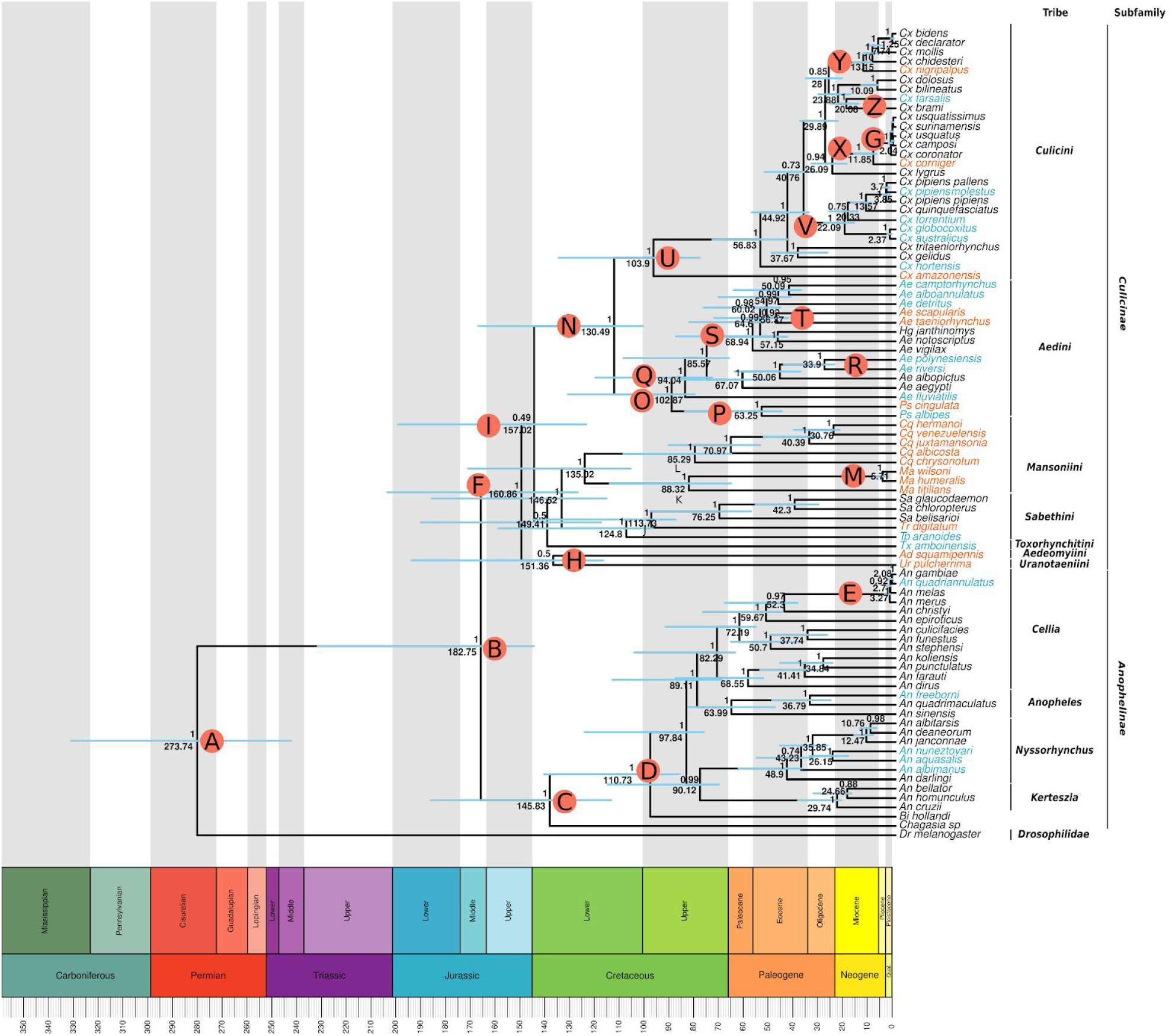
Evolutionary timescale of Culicidae family. Bayesian three reconstructed using nucleotide sequences of PCGs partitioned by gene and codon positions. Blue bars represent the HPD95%. Numbers above and below the bars show the posterior probability and the predicted median dating for each node. Specific words inside the circles represent the nodes discussed in the text.

The subfamily *Culicinae* formed a monophyletic group with the TMRCA (The most recent common ancestral) occurring around 160 MYA in Jurassic period (Figure 3, node F). The basal lineage from Culicinae subfamily was two species representing the *Uranotaenia* and *Aedeomyia* genera which shared a common ancestor around 151 MYA (Figure 4, node H). The split of *Culicini* and *Aedini* tribes from *Toxorhynchitini, Sabethini, Mansoniini* has occurred around 157 MYA (Figure 4, node I). The *Toxorhynchites* genus has shown to be basal in relation to *Sabethini* and *Mansoniini* tribes. Among *Sabethini* members, the *Tripteroides* genus was positioned as a basal lineage and *Trichoprosopon* genus speciated from other *Sabethes* species around 113 MYA (Figure 4, node J). The *Mansoniini* tribe has been placed as a sister clade to the *Sabethini* tribe. The *Mansonia* and *Coquillettidia* genera were both monophyletic with speciation processes starting around 88 and 85 MYA respectively (Figure 4, nodes K and L, respectively). The *Ma. titillans* species showed basal in relationship to *Ma. wilsoni* and *Ma. humeralis* that speciated in Neogene period (5,71 MYA, node M in Figure 4). Inside the *Coquillettidia* genus, *Cq. chrysonotum* showed a basal position in relationship the other species. The diversification inside *Coquillettidia* occurred between 85 to 30 MYA with origin of *Cq. chrysonotum, Cq. albicosta, Cq. juxtamansonia* and *Cq. venezuelensis* + *Cq. hermanoi* respectively (Figure 4). The diversification between *Culex* and *Aedini* taxa occurred in the Cretaceous period around 130 MYA (Figure 4, node N). While the split of *Aedes* and *Psorophora* genera occurred around 102 MYA and the speciation of *Ps. albipes* and *Ps. cingulata* occurred in Paleogene around 63 MYA (Figure 4, node O and P, respectively). Among the *Aedes* species, *Ae. fluviatilis* is the basal and early diverged species (94 MYA, node Q in Figure 4) from the genus. *Ae. polynesiensis* and *Ae. riversi* were closely to *Ae, albopictus* (Figure 4, node R). Another clade formed closely to *Ae. aegypti* clade was composed by species from *Ochlerotatus* (*Ae. vigilax, Ae. taeniorhynchus, Ae. scapularis, Ae. detritus* and *Ae. camptorhynchus*), *Finlaya* subgenera (*Ae. notoscriptus* and *Ae. alboannulatus*) and *Haemagogus* genus, where *Ae. vigilax* was the basal species (Figure 4, node S). The *Finlaya* subgenus has a paraphyletic status when the positioning of *Ae. alboannulatus* and *Ae. notoscriptus* is observed (Figure 4). The neotropical species *Ae. taeniorhynchus* and *Ae. scapularis* formed a clade and diverged between themself around 56 MYA (Figure 4, node T). Among *Culex* species, *Cx. amazonensis* showed to be the basal and the earlier diverged species from the genus with the split from the other species occurring around 103 MYA (Figure 4, node U). The *pipiens* group showed an origin around 22 MYA where the Australian species *Cx. australicus* and *Cx. globocoxitus* have been placed in basal position in relationship to other *Cx. pipiens* species (Figure 4, node V). The *Cx. torrentium* was placed inside *pipiens* group. The *Cx. corniger* was placed as basal species from *coronator* group of *Culex* with speciation around 11 MYA as well as *Cx. nigripalpus* that speciated from *Cx. chidesteri, Cx. mollis, Cx. declarator* and *Cx. bidens* around 13 MYA (Figure 4, node X and Y respectively). While *Cx. tarsalis* formed a clade with *Cx. brami* with a speciation time of 20 MYA (Figure 4, node Z).

**Figure 4.**
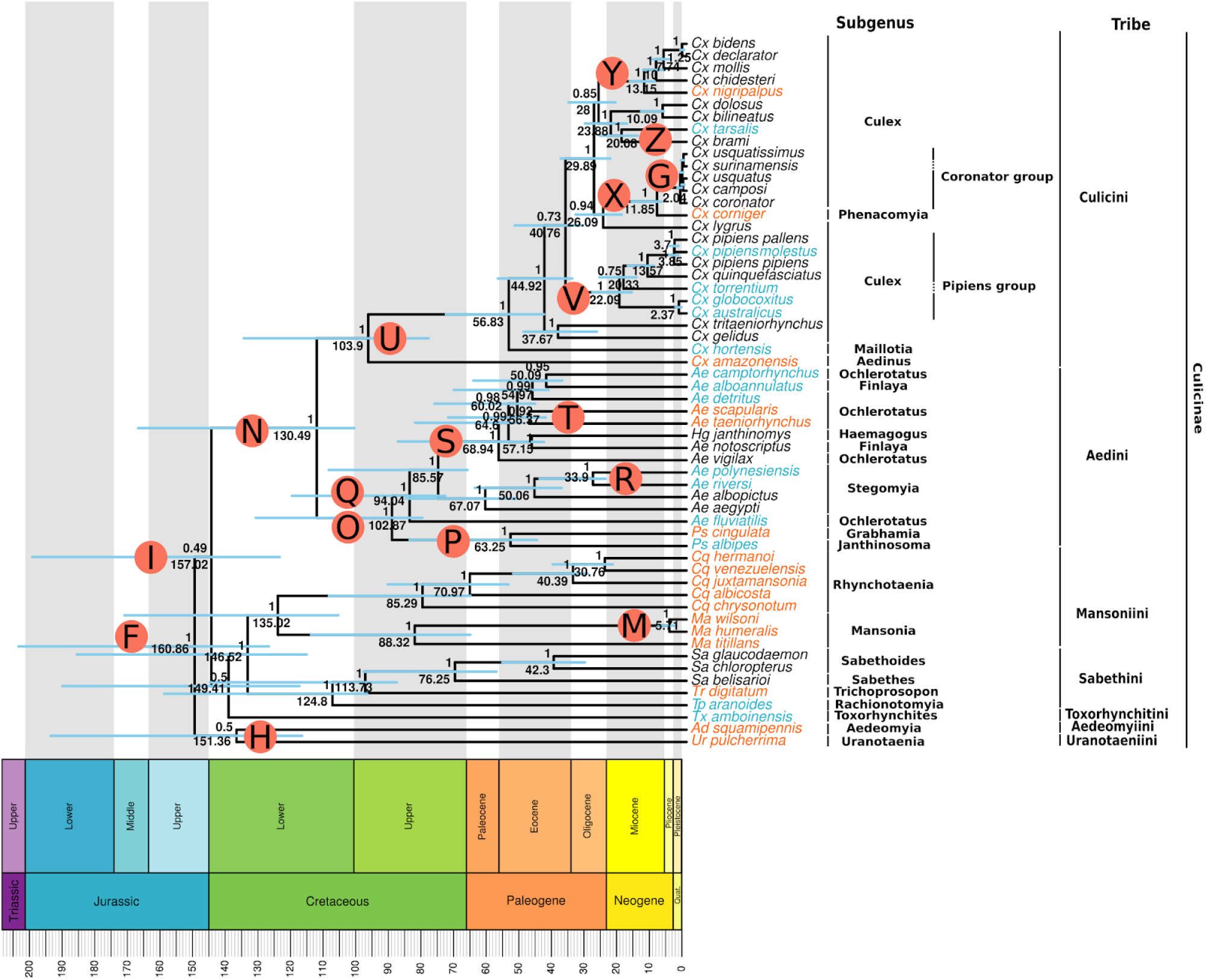
Evolutionary timescale of Culicinae subfamily. Tree was generated from BEAST analysis of partitioned PCG taking into account the split of codon positions. Blue bars in the nodes represent the HPD95%. The numbers above and below the bars show the posterior probability and the predicted median dating respectively for each node. Specific words inside the circles represent the nodes discussed in the text.

### Lineage through time (LTT) section

The LTT analysis showed the radiation processes within Culicidae family were constant through the last million years with some peaks (Figure 5A). After the first lineage emergences splitting the subfamilies *Anophelinae* and *Culicinae* occurred a rise in speciation processes up to 100 MYA in Cretaceous period. Another speciation peak occurred close to 75 MYA. After the peak close to 75 MYA the speciation processes occured in a more constant pace until before 25 MYA and showed a rise in speciation only in the last million years. Within *Anophelinae* subfamily a constant pattern of speciation occurred with few peaks (Figure 5B). Likewise, the LTT analysis within *Culicinae* subfamily showed a similar pattern seen in *Anophelinae* LTT (Figure 5C).

**Figure 5.**
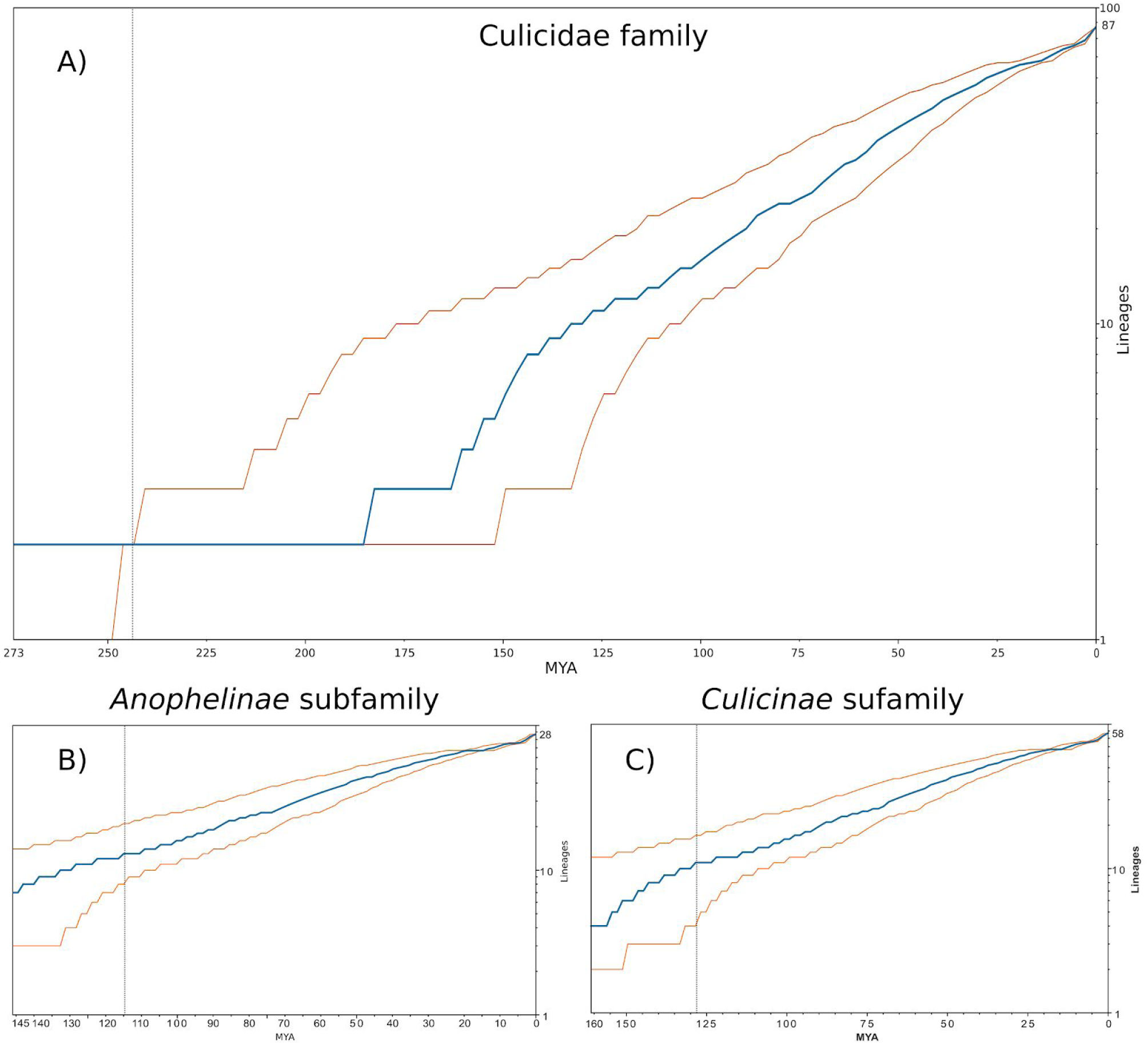
Lineages through time (LTT) plots representing the radiation within each taxa. A) LTT analysis showing the radiation within Culicidae family. B) LTT analysis showing the radiation within *Anophelinae* subfamily. C) LTT analysis showing the radiation within *Culicinae* subfamily. X-axis represents the time in million years ago (MYA). Y-axis shows the number of lineages. Blue line represents the LTT trace and orange lines show the confidence intervals.

## Discussion

Mitogenomes have been widely used to elucidate the evolutionary history of several species of animals and plants and can also be used as barcode sequences for species identification ^14,55,56^. The first mosquito mitogenome from *Anopheles gambiae* was sequenced using the Sanger method on PCR amplified fragments ^25^ and further mitochondrial genomes were slowly sequenced along with whole genome projects. Currently, most studies have been using high throughput sequencing or PCR amplification followed by high throughput sequencing to characterize several mitogenomes at once ^20,21,24^ in a wide range of insect species showing promising results to reconstruct mitogenomes ^30,57^ Here, we performed low-coverage sequencing to assemble and characterize the mitogenomes from many mosquito species generating around 4.95 million reads per species. We were able to assemble sixteen draft mitogenomes from Culicidae species, not characterized so far, belonging to 8 different genera. Richter et al. (2015)^30^ and collaborators suggested that a minimum of 10 million reads are needed to recovery mitogenomes with higher coverage breadth and datasets having around 1 million reads usually generate highly incomplete mitogenomes, but we were able to assemble nearly complete mitogenomes with as low as 1.1 million reads. Besides, the draft genomes assembled contained enough phylogenetic markers necessary for robust phylogenetic analysis.

In addition, we also reconstructed mitochondrial genomes from available RNA-Seq data. We were able to reconstruct twenty additional nearly complete draft mitogenomes for Culicidae species not studied so far at the mitochondrial genomic level. No study has been able to reconstruct complete mitochondrial genomes from RNA-Seq data, mainly due the endonuclease activity on transcripts or loss of mitochondrial transcripts due to the enrichment steps normally used during the sequencing library construction ^28^. However, the remaining mitochondrial data available in different RNA-Seq datasets still can be used to mine mitogenome sequences for understudied species ^58–60^. The datasets used for mitogenomes characterization showed around 0.073 to 8.034% of mitochondrial reads. In total, we obtained 37 draft mitogenomes representing 11 genera (*Anopheles, Uranotaenia, Aedeomyia, Toxorhynchites, Tripteroides, Trichoprosopon, Mansonia, Coquillettidia, Psorophora, Aedes* and *Culex*).

Several efforts have been made to better understand the taxonomic status inside of the Culicidae family, but most studies that included a substantial number of species employed only morphological data ^61^ and the ones using molecular information are mainly based on a low number of species or used limited molecular markers ^62–65^. Hence, there are still many non-studied species and unresolved phylogenetic relationships in genera such as *Aedes, Armigeres, Coquillettidia, Culex, Mansonia, Mimomyia, Psorophora, Topomyia, Tripteroides, Toxorhynchites, Uranotaenia* and *Wyeomyia* ^61^. The *Anophelinae* subfamily phylogenetic relationship was recently investigated using the mitochondrial genomes from many species and the authors proposed a number of taxonomic status changes such as the elevation of some groups from the subgenus to genus level ^19^. Some other studies also characterized the mitogenomes of some species to solve the phylogenetic positioning of culicids, however, most of them focused on a specific group (*Anopheles*, *Culex* and *Sabethes*) or few genera of the Culicidae family^10,11, 19–21,66^.

The phylogenetic analysis including the thirty-five mitogenomes assembled in this study comprising eight Culicidae genera is highly congruent regarding the monophyly of large species groups. Both Culicinae and Anophelinae subfamily and *Anopheles, Sabethes*, *Mansonia,* Coquillettidia, *Psorophora* and *Culex* genera were monophyletic. Moreover, we also observed similar dating, estimates as reported in the literature, for some key ancestors. For instance, the split between Culicidae and other dipterans occurred around 273 MYA (HPD95%:243.79-332.41) while other studies suggested that this split could have occurred around 259 and 260 MYA using mitogenomes and phylogenomics analysis respectively ^11,65^. The split between the two subfamilies: *Anophelinae* and *Culicinae* occurred in the Jurassic period around 182 MYA (HPD95%:145.88-232.95). Similar estimates were obtained in other studies around 190-195 MYA ^67,68^. Different evolutionary rate of molecular markers, incomplete species sampling and different algorithms used to reconstruct the species phylogeny could result in different time estimates (Hao et al. (2017)).

The evolutionary history of the *Anophelinae* subfamily has been more extensively studied considering the number of species analyzed and the different morphological and molecular markers used including whole phylogenomic analysis ^64^ as well as mitogenomes (FOSTER et al. 2017). However, these studies have not assessed the diversification of some basal groups such as *Chagasia* and *Bironella*. In our analysis, those groups showed to be the early diverged lineages from *Anophelinae* subfamily, emerging in the Upper and Lower Cretaceous, respectively (Figure 3). The *Bironella* genus showed as an ancestral lineage in relationship to the *Anopheles* genus including all subgenera assessed in our analysis such as *Kerteszia*, *Nyssorhynchus, Anopheles* and *Cellia*. This contrasts with other findings that positioned this genus inside the *Anopheles* genus using mitochondrial protein sequences from mitogenomes ^19^. Previous studies, using both nuclear ribosomal sequences and fragments of mitochondrial genes COI and COII of *Bi. Gracilis* ^69^ and *Bi. hollandi* ^70^, have already suggested that positioning of *Bironella* inside *Anopheles* genus. These contrasting results show that the *Bironella* genus position and phyletic status is still open and a wide sampling of the genus is needed to solve it. Regarding the *Anopheles* species our analysis using mitogenomes showed a similar positioning as seen in other studies ^70,71^.

The radiation inside the *Culicinae* subfamily is older than *Anophelinae* around 160 MYA (HPD95%:128.09-204.91) in the Jurassic period (Supplementary figure 4 and 5). Inside Culicinae subfamily the clade composed by *Uranotaenia* and *Aedeomyia* genera are the earliest diverging branch with the split from the other genera occurring around 151 MYA (HPD95%:117.92-195.06) (Figure 4). The *Toxorhynchites amboinensis* species analyzed was placed as a sister group of *Sabethini* and *Mansoniini* tribes showing a very ancient split from these genera around 149 MY (HPD95%:118.74-191.45) corroborating findings from studies based on morphological characters which suggested that *Aedeomyia, Uranotaenia* and *Toxorhynchites* genera are ancient and basal groups inside of the Culicinae subfamily ^62,72,73,74^. However, two studies based on six nuclear genes and 18S rDNA have shown the positioning of *Uranotaenia sapphirina* more closely related to *Culicini* and *Aedini* tribes respectively ^63,75^ which warrants more in depth analysis in order to establish the true genus positioning and phyletic status.

Regarding the *Sabethini* tribe, our results are in line with previous works showing the monophyly of tribe, the basal positioning of *Tripteroides* (*Tp. aranoides*) and the sister positioning of *Trichoprosopon* genus (*Tr. digitatum*) ^20,24,63^ (Figure 4). Our analysis demonstrated the origin of *Sabethini* tribe around 124 MYA (HPD95%:97.93-160.21) in the Lower cretaceous period. Although limited data exists about the arboviral vectorial capacity of different mosquito species of this tribe, our phylogenetic data for the *Sabethini* group associated with the competence of different New World *Sabethes* species to transmit *Yellow fever virus* (YFV) ^76–78^ and the demonstrated absence of competence of Old World *Tripteroides* species ^79^ suggests that arboviral transmission competence emerged in the New World in the ancestral of the *Sabethes* genus between Upper Cretaceous and Paleogene (76.25 MYA - HPD95%:58.26-99.06) after the split from the *Tripteroides* genus.

Up to now, few studies have investigated the phylogenetic positioning and speciation time of the neotropical *Mansoniini* tribe. In our analysis, the eight species from the *Mansoniini* tribe formed a monophyletic group and are a sister group of the *Sabethini* tribe. Our results are in contrast with Reidenbach et al. (2009) analysis that positioned a single *Coquillettidia* species (*Cq. pertubans*) as a sister group of *Aedini.* Our dataset covers a larger number of species from the *Mansoniini* tribe and Reidenbach et al. (2009) did not include any *Mansonia* species which probably biased their analysis. Both *Coquillettidia* and *Mansonia* genera are monophyletic groups (Figure 4). Although vector capacity information for most of the *Mansoniini* species investigated here are unknown, some studies were able to detect arboviruses such as Mayaro virus and Saint Louis encephalitis virus in *Cq. venezuelensis* and *Ma. titillans* respectively ^80–82^. Furthermore, *Ma. titillans* species showed a vector competence to Venezuelan Equine Encephalomyelitis Virus being considered as a second vector of this arbovirus in the Peruvian Amazon ^83^. Therefore, the basal positioning of these two species inside of the *Mansonia* and *Coquillettidia* genera suggest that others *Mansoniini* species may also be competent for transmitting arboviruses.

Within the *Aedini* tribe our results have shown the same basal positioning of *Psorophora* genus seen by Reidenbach et al. (2009). However, we estimated the split of *Janthinosoma* e *Grabhamia* subgenera through the first sequenced mitogenomes of *Psorophora* genus, *Ps. cingulata and Ps. albipes,* occurring around 63 MYA (HPD95%:45.84-84.94) on Paleogene period. Regarding the *Aedes* genus, our results showed a paraphyletic group encompassing a single species from the *Haemagogus* genus which corroborates other findings with a larger number of *Haemagogus* species ^84^. Besides, paraphyletic groups was also observed for *Ochlerotatus* (*Ae. fluviatilis, Ae. taeniorhynchus*, *Ae. scapularis, Ae. vigilax, Ae. detritus* e *Ae. camptorhynchus*) and *Finlaya* (*Ae. notoscriptus* e *Ae. alboannulatus*) subgenera while *Stegomyia* (*Ae. aegypti, Ae. albopictus, Ae. riversi, Ae. polynesiensis*) subgenera formed a monophyletic group (Figure 4). A previous study based on morphological cladistic analysis suggested the monophyly of *Ochlerotatus* and *Finlaya* subgenera ^85^. Inside of the *Aedes* genus, *Ae. fluviatilis* is the earliest branch split around 94 MYA (HPD95%:74.02-121.02), in contrast to other study that positioned it within other *Aedes* branches ^84^. In summary, the *Aedes* genus is a paraphyletic group showing several phylogenetic incongruences even among studies that used different markers and species representatives. Hence further reclassification is needed following the current knowledge of phylogenetic relationships of these species.

Some studies demonstrated the vectorial competence of *Ae. fluviatilis* for Dengue Virus (DENV) in experimental conditions ^86,87^. Besides, this species is also a potential vector for *Dirofilaria immitis* transmission ^88^. The basal positioning of *Ae. fluviatilis* in relation to the other *Aedes* species suggests that the vectorial capacity for arboviruses and filarial transmission is an ancestral trait of the *Aedes* genus. This trait likely emerged between the Cretaceous and Paleogene periods (94.04-33.9 MYA - HPD95%:74.02-121.02/24.82-45.09). It is corroborated by the abundant knowledge about more recent divergent species that inherited this trait such as *Ae. aegypti* and *Ae. albopictus*, that are known vectors of DENV, YFV and *D. immitis* ^3,4,89,90^. Moreover, other species such as *Ae. polynesiensis* ^91,92^*, Ae. taeniorhynchus* ^93^*, Ae. vigilax* ^94^*, Ae. scapularis* ^93^, *Ae. notoscriptus* ^94^ are reported as potential or confirmed vectors of filarial worms.

Regarding the *Culex* genus, our analysis showed that *Cx. amazonensis* and *Cx, hortensis* are one of the earliest diverged species from this genus dating around 103 MYA (HPD95%:79.07-135.87) and 56 MYA (HPD95%:43.69-73.86), respectively. Our results are in agreement with ^95^ cladistic morphological analysis concerning the basal positioning of those species, however our mitogenomics data support *Cx. amazonensis* as the earliest divergent species instead of *Cx. hortensis*. Our analysis have placed *Cx. nigripalpus* as sister group of clade composed by *Cx. chidesteri, Cx. mollis, Cx. declarator* e *Cx. bidens* with radiation around 13 MYA (HPD95%:9.82-17.37). While, *Cx. corniger* has placed as sister lineage from *Coronator* group. A previous study using a fragment of the COI gene have already suggest this positioning ^96^ and our mitogenomics analysis supported this placement showing it speciation time around 11.85 MYA (HPD95%:7.78-16.85) in the Neogene period. Much have been discussed about the members of *Pipiens* group, if *Cx. pipiens* consist in a species or a group of sibling species ^95^. Some authors describe the *Pipiens* group harboring the following species: *Cx. pipiens pipiens, Cx. quinquefasciatus, Cx. pipiens pallens, Cx. pipiens molestus, Cx. australicus* e *Cx. globocoxitus* ^95,97^. Other similar species such as *Cx. torrentium* have not been considered as a member of *Pipiens* group due its genetic divergence to other species of the group ^98^. A study based on analysis of ITS1 and ITS2 has already demonstrated the close relationship of *Cx. torrentium* with *Pipiens* group ^99^. Our analysis have positioned *Cx. torrentium* within *Pipiens* group with australian members *Cx. globocoxitus* and *Cx. australicus* as basal clade suggesting that *Cx. torrentium* may be a true species from the *Pipens* group. Our dating analysis estimated that the *Pipiens* group split from other *Culex* species around 40 MYA (HPD95%:31.79-52.73) and the speciation process inside of *Pipiens* group occurred from 22 MYA (HPD95%:16.73-28.91) up to 2.37 MYA (HPD95%:1.53-3.37). Although the lower divergence time among some members of *Pipiens* group each “species” has specific ecological, physiological and behavioral characteristics ^98,100^.

Our LTT analysis showed that the diversification within Culicidae family occured in a more or less continuous processes over time. However, at some points occurred a rise in speciation processes. Tang et al. (2018) described that at least three shifts in speciation rates in Culicidae family occurred around 180-195 MYA and the last around 30-24 MYA. This shift on speciation rates demonstrated that they are correlated with the split of subfamilies and the first lineages of mosquitoes. The latest shifts in recent speciation (30-24 MYA) correlate with some *Anopheles* and *Culex* speciation as seen in our results. However, it is important to note that the number of species and the representation of the species used in the phylogenetic analysis can drastically affect the LTT plot shape. Therefore, it is important to see those results with care to interpret speciation rate in mosquitoes.

## Conclusion

Overall, we characterized the phylogenetic position and speciation time of the main groups of Culicidae family which emerged in the last 182 MYA between the Jurassic and Paleogene periods. Most of the different genera emerged in this range of time, but some recent speciation occurred as in the *Culex* genus. Interestingly, a burst in mammals speciation also occurred in the Neogene period likely driving the speciation of these species at that time ^67,101^. Furthermore, the new phylogenetic knowledge generated allowed us to propose new hypotheses about some mosquito traits emergence and maintenance related with vector competence. More in depth studies trying to tease apart these different mechanisms considering the phylogeny of the Culicidae tree will benefit from the information generated in this work.

## Acknowledgements

This research was supported by Fundação Oswaldo Cruz and Conselho Nacional de Desenvolvimento Científico e Tecnológico (CNPq) under the project number PROEP/IAM (400742/2019-5). da Silva, A.F received a masters scholarship from Coordenação de Aperfeiçoamento de Pessoal de Nível Superior (CAPES). We thank Cláudio Júlio da Silva and Edivaldo José Apolinário from Núcleo de Vigilância à Saúde e do Meio Ambiente by the support in mosquito sampling from Moreno municipality. We thank the Núcleo de Bioinformática and Núcleo de Plataformas Tecnológicas (NPT) of IAM by the computational infrastructure and sequencing platform respectively.

## Author contributions statement

Wallau, G.L conceived, designed the research, performed mosquito sampling and wrote the paper. da Silva, A.F performed the mosquito sampling, molecular experiments, bioinformatics analysis and drafted the paper. Machado, L.C performed the mosquito sampling, molecular experiments and contributed to drafted the manuscript. Paula, M.B and Pessoa, C.J.S performed the taxonomic identification and contributed to write the paper. Bronzoni, R.V.M and Santos, M.A.V.M performed mosquito sampling and contributed to write the paper. All authors read and approved the final version of the manuscript.

---

**Supplementary table 1.**
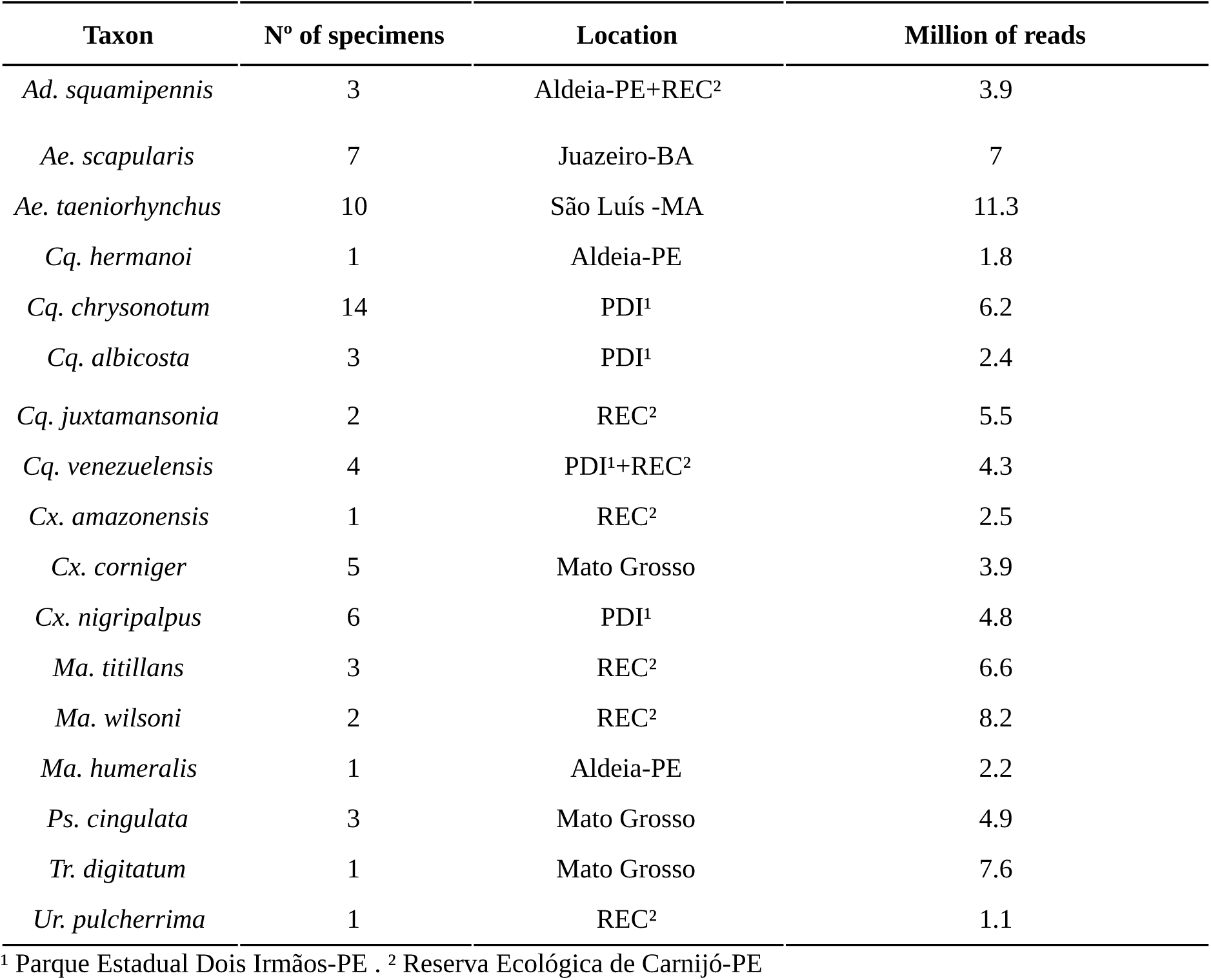
Information of sequenced samples.

**Supplementary table 2.**
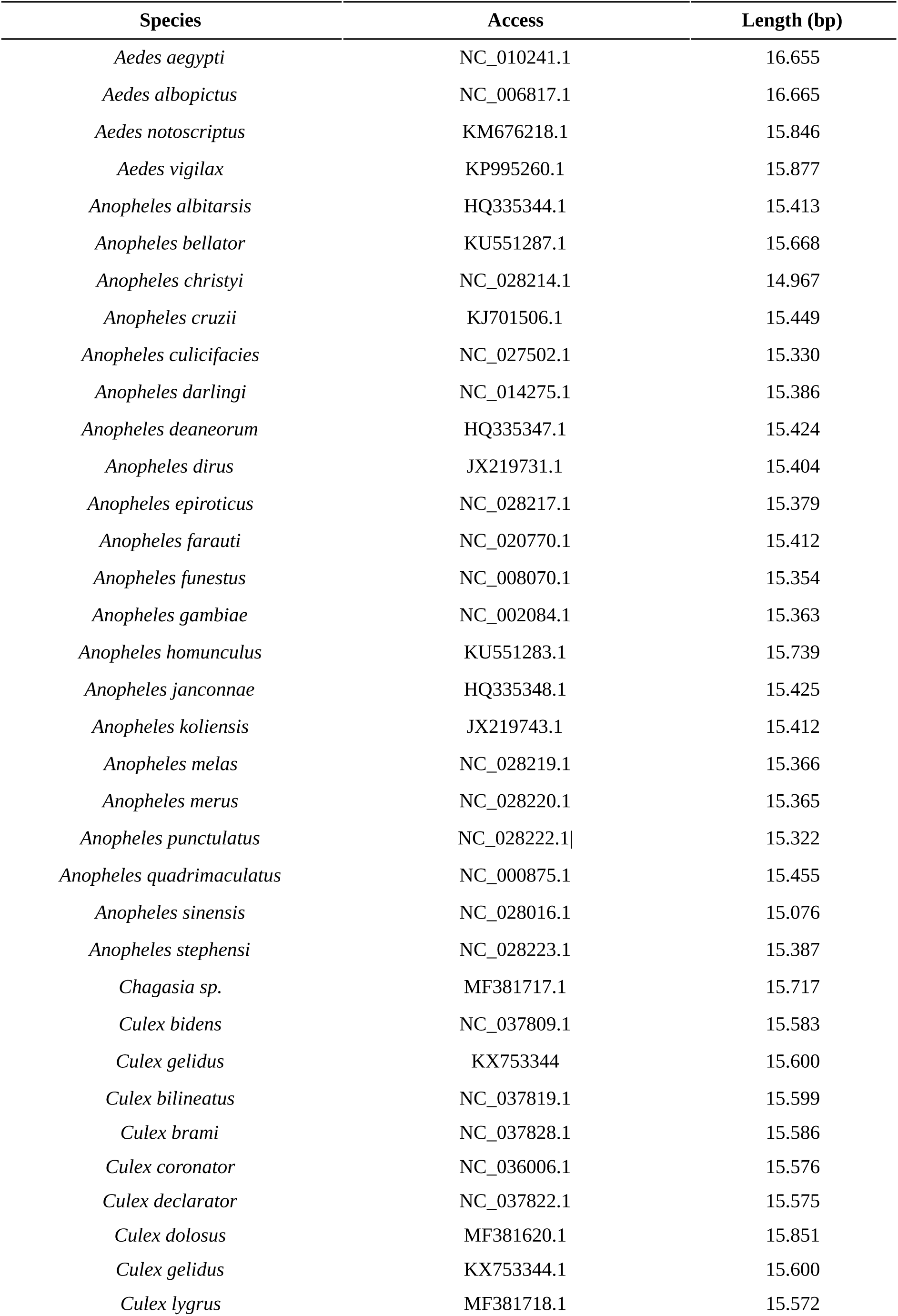
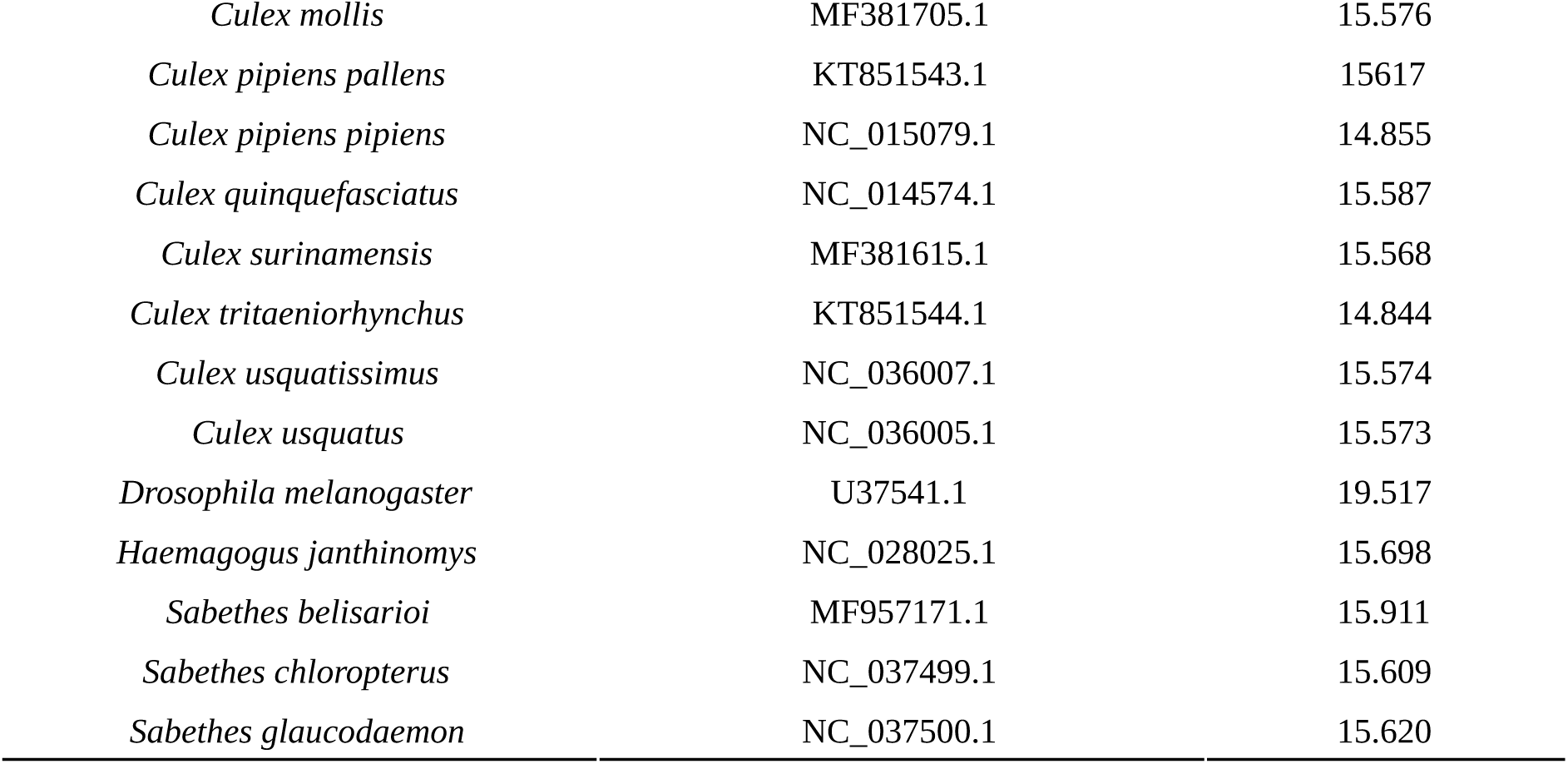
Access list of mitochondrial genomes recovered from NCBI.

**Supplementary table 3.**
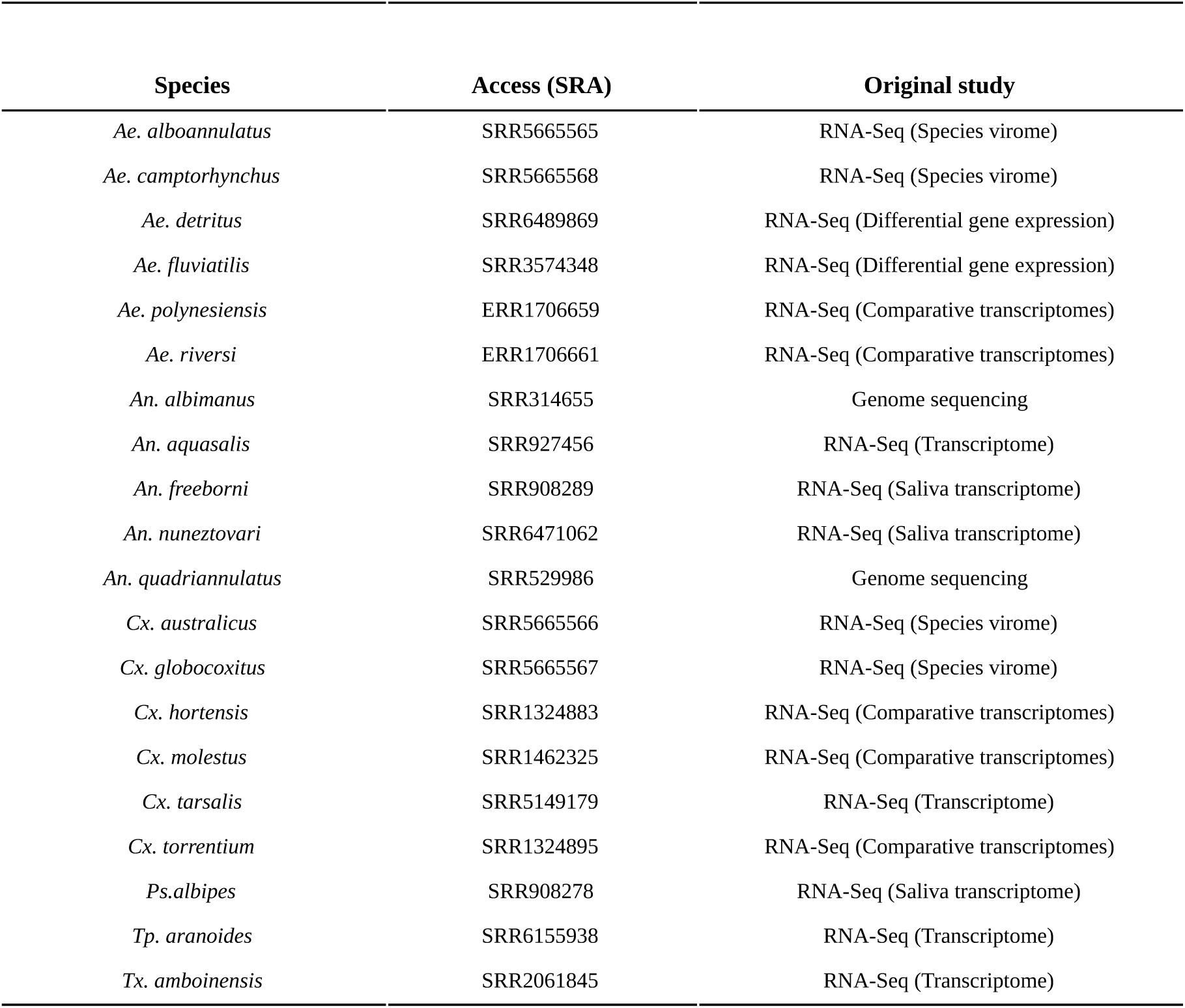
Information about minered datasets from NCBI SRA database.

**Supplementary table 4.**
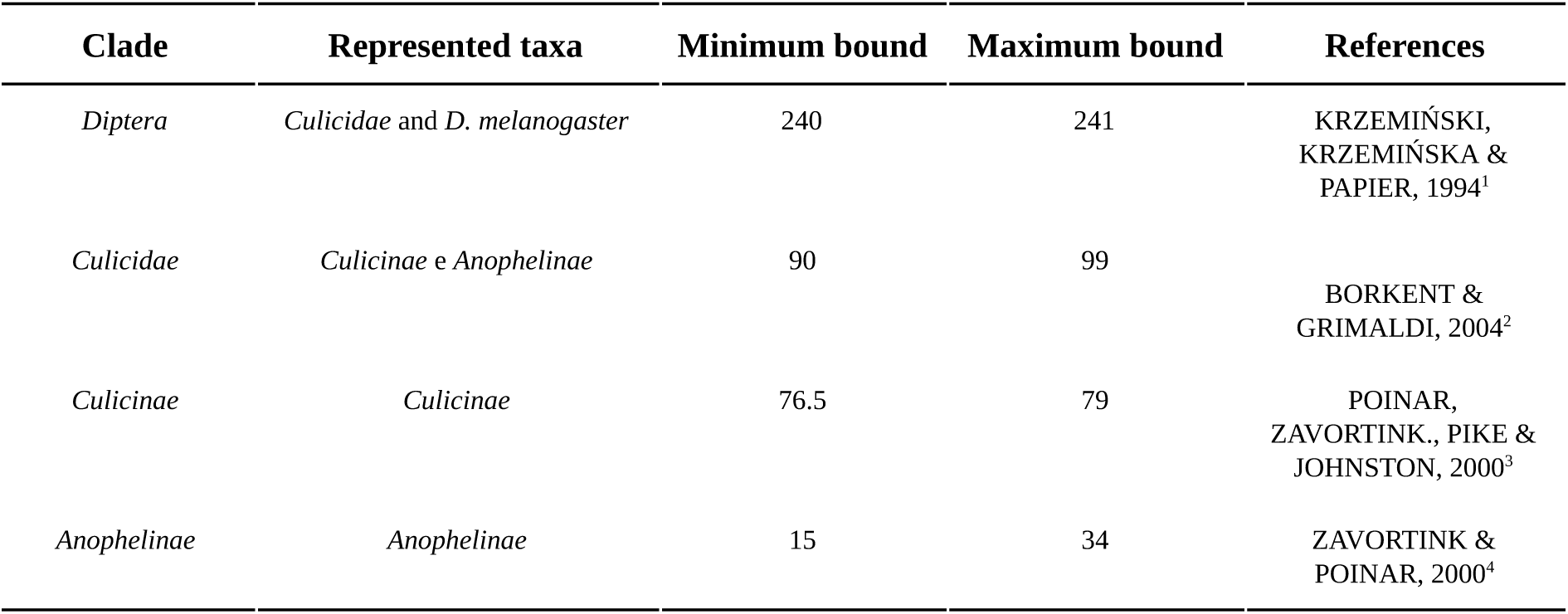
Calibration points used in BEAST 1.8.4

**Supplementary table 5.** Evolutionary models used in BEAST analysis.

| Partition | Best nucleotide model <sup>1</sup> | Best protein model <sup>2</sup> |
| --- | --- | --- |
| Nucleotide mitogenome alignment | GTR+G+I | - |
| Concatenated protein alignment | - | mtREV+G+I |
| Atp6 | GTR+G+I | mtREV+G+I |
| Atp8 | GTR+G | JTT+G+I |
| COIII | GTR+G+I | mtREV+G+I |
| COII | GTR+G+I | mtREV+G |
| COI | GTR+G+I | JTT+G+I |
| CytB | GTR +G+I | mtREV+G+I |
| Nad1 | GTR +G+I | JTT+G+I |
| Nad2 | GTR +G+I | JTT+G |
| Nad3 | GTR +G+I | mtREV+G+I |
| Nad4L | GTR +G+I | mtREV+G |
| Nad4 | GTR +G+I | JTT+G+I |
| Nad5 | GTR +G+I | JTT+G |
| Nad6 | GTR +G+I | mtREV+G |
| rrnL | GTR +G+I | - |
| rrnS | GTR +G+I | - |

<sup>1</sup>Best model as suggested by SMS. <sup>2</sup> More likely model suggested by Prottest and present in BEAST 1.8.4.

**Supplementary table 6.** Annotation of draft mitogenomes using MITOS.

| Species | *nad2* | *COI* | *COII* | *atp8* | *atp6* | *COIII* | *nad3* | *nad5* | *nad4* | *nad4l* | *nad6* | *cytb* | *nad1* | *rrnL* | *rrnS* | *tRNA* | *PCG* | *rRNA* | Total of genes |
| --- | --- | --- | --- | --- | --- | --- | --- | --- | --- | --- | --- | --- | --- | --- | --- | --- | --- | --- | --- |
| ***Ad. squamipennis*** | 1 | 1 | 1 | - | - | 1 | 1 | 1 | 1 | - | - | 1 | 1 | 1 | 1 | 16 | 9 | 2 | 27 |
| *Ae. alboannulatus* | 1 | 1 | 1 | - | - | 1 | 1 | 1 | 1 | 1 | 1 | 1 | 1 | 1 | 1 | 13 | 11 | 2 | 26 |
| *Ae. camptorhynchus* | 1 | 1 | 1 | 1 | - | 1 | 1 | 1 | 1 | 1 | 1 | 1 | 1 | 1 | 1 | 16 | 12 | 2 | 30 |
| *Ae. detritus* | 1 | 1 | 1 | 1 | 1 | 1 | 1 | 1 | 1 | 1 | 1 | 1 | 1 | 1 | 1 | 16 | 13 | 2 | 31 |
| *Ae. fluviatilis* | 1 | 1 | 1 | 1 | 1 | 1 | - | 1 | 1 | 1 | 1 | 1 | 1 | 1 | - | 4 | 12 | 1 | 17 |
| *Ae. polynesiensis* | 1 | 1 | 1 | 1 | - | 1 | 1 | 1 | 1 | 1 | 1 | 1 | 1 | 1 | 1 | 20 | 12 | 2 | 34 |
| *Ae. riversi* | - | 1 | 1 | 1 | 1 | 1 | 1 | 1 | 1 | - | - | 1 | 1 | 1 | - | 6 | 10 | 1 | 17 |
| ***Ae. scapularis*** | 1 | 1 | 1 | 1 | 1 | 1 | - | 1 | 1 | - | 1 | 1 | 1 | - | 1 | 13 | 11 | 1 | 25 |
| ***Ae. taeniorhynchus*** | 1 | 1 | 1 | 1 | 1 | 1 | 1 | 1 | 1 | 1 | 1 | 1 | 1 | 1 | 1 | 22 | 13 | 2 | 37 |
| *An. albimanus* | 1 | 1 | 1 | 1 | - | 1 | 1 | 1 | 1 | 1 | 1 | 1 | 1 | 1 | 1 | 20 | 12 | 2 | 34 |
| *An. aquasalis* | 1 | 1 | 1 | 1 | - | 1 | - | 1 | 1 | - | - | 1 | 1 | 1 | 1 | 3 | 9 | 2 | 14 |
| *An. freeborni* | 1 | 1 | 1 | 1 | - | 1 | 1 | 1 | 1 | 1 | 1 | 1 | 1 | 1 | 1 | 19 | 12 | 2 | 33 |
| *An. nuneztovari* | 1 | 1 | 1 | 1 | - | 1 | 1 | 1 | 1 | 1 | 1 | 1 | 1 | 1 | 1 | 9 | 12 | 2 | 23 |
| *An. quadriannulatus* | 1 | 1 | 1 | 1 | - | 1 | 1 | 1 | 1 | 1 | 1 | 1 | 1 | 1 | 1 | 20 | 12 | 2 | 34 |
| ***Cq. albicosta*** | 1 | 1 | 1 | 1 | 1 | 1 | 1 | 1 | 1 | 1 | 1 | 1 | 1 | 1 | 1 | 21 | 13 | 2 | 36 |
| ***Cq. chrysonotum*** | 1 | 1 | 1 | - | 1 | 1 | 1 | 1 | 1 | 1 | 1 | 1 | 1 | 1 | 1 | 20 | 12 | 2 | 34 |
| ***Cq. hermanoi*** | 1 | 1 | 1 | 1 | 1 | 1 | 1 | 1 | 1 | - | - | 1 | - | 1 | 1 | 14 | 10 | 2 | 26 |
| ***Cq. juxtamansonia*** | 1 | 1 | 1 | - | 1 | 1 | - | 1 | 1 | - | - | 1 | - | 1 | 1 | 8 | 8 | 2 | 18 |
| ***Cq. venezuelensis*** | 1 | 1 | 1 | 1 | 1 | 1 | 1 | 1 | 1 | 1 | 1 | 1 | 1 | 1 | 1 | 19 | 13 | 2 | 34 |
| ***Cx. amazonensis*** | 1 | 1 | 1 | 1 | 1 | 1 | 1 | 1 | 1 | 1 | 1 | 1 | 1 | 1 | 1 | 21 | 13 | 2 | 36 |
| *Cx. australicus* | 1 | 1 | 1 | 1 | 1 | 1 | 1 | 1 | 1 | 1 | 1 | 1 | 1 | 1 | 1 | 17 | 13 | 2 | 32 |
| ***Cx. corniger*** | 1 | 1 | 1 | 1 | - | 1 | - | 1 | 1 | 1 | - | 1 | - | 1 | 1 | 9 | 9 | 2 | 20 |
| *Cx. globocoxitus* | 1 | 1 | 1 | 1 | - | 1 | 1 | 1 | 1 | 1 | 1 | 1 | 1 | 1 | 1 | 17 | 12 | 2 | 31 |
| *Cx. hortensis* | 1 | 1 | 1 | - | - | 1 | 1 | 1 | 1 | 1 | 1 | 1 | 1 | 1 | 1 | 12 | 11 | 2 | 25 |
| *Cx. molestus* | 1 | 1 | 1 | 1 | 1 | 1 | 1 | 1 | 1 | 1 | 1 | 1 | 1 | 1 | 1 | 6 | 13 | 2 | 21 |
| ***Cx. nigripalpus*** | 1 | 1 | 1 | 1 | 1 | 1 | 1 | 1 | 1 | 1 | 1 | 1 | 1 | 1 | 1 | 22 | 13 | 2 | 37 |
| *Cx. tarsalis* | 1 | 1 | 1 | 1 | - | 1 | 1 | 1 | 1 | 1 | 1 | 1 | 1 | 1 | 1 | 19 | 12 | 2 | 33 |
| *Cx. torrentium* | 1 | 1 | 1 | 1 | 1 | 1 | 1 | 1 | 1 | 1 | 1 | 1 | 1 | 1 | 1 | 11 | 13 | 2 | 26 |
| ***Ma. wilsoni*** | 1 | 1 | 1 | 1 | 1 | 1 | 1 | 1 | 1 | 1 | - | 1 | 1 | 1 | 1 | 14 | 12 | 2 | 28 |
| ***Ma. humeralis*** | - | 1 | 1 | - | 1 | - | 1 | 1 | - | - | - | 1 | 1 | 1 | 1 | 5 | 7 | 2 | 14 |
| ***Ma. titillans*** | 1 | 1 | 1 | - | 1 | 1 | 1 | 1 | 1 | - | 1 | 1 | 1 | 1 | 1 | 14 | 11 | 2 | 27 |
| ***Ps. cingulata*** | 1 | 1 | 1 | 1 | 1 | 1 | 1 | 1 | 1 | 1 | 1 | 1 | 1 | 1 | 1 | 21 | 13 | 2 | 36 |
| *Ps. albipes* | 1 | 1 | 1 | 1 | - | 1 | 1 | 1 | 1 | 1 | 1 | 1 | 1 | 1 | 1 | 19 | 12 | 2 | 33 |
| *Tr. aranoides* | 1 | 1 | - | - | - | 1 | - | 1 | - | 1 | - | 1 | 1 | 1 | 1 | - | 7 | 2 | 9 |
| ***Tr. digitatum*** | 1 | 1 | 1 | - | 1 | 1 | 1 | 1 | 1 | 1 | - | 1 | - | 1 | 1 | 16 | 10 | 2 | 28 |
| *Tx. amboinensis* | 1 | 1 | 1 | 1 | 1 | 1 | 1 | 1 | 1 | 1 | 1 | 1 | 1 | 1 | 1 | 19 | 13 | 2 | 34 |
| ***Ur. pulcherrima*** | 1 | 1 | 1 | - | 1 | 1 | 1 | 1 | 1 | 1 | 1 | 1 | 1 | 1 | 1 | 18 | 12 | 2 | 32 |

Legend: Bold names represent sequenced mitogenomes. Disrupted or duplicated gene are represented by bold numbers. Unidentified gene are represented by -.

**Supplementary table 7.** Predicted radiation values from Bayesian inference analysis for studied species. The HPD95% values represents the confidence intervals for radiation between compared taxa.

| Node<sup>1</sup> | HPD 95% | Median dating | Posterior probability |
| --- | --- | --- | --- |
| A | 243.79-332.41 | 273.74 | 1 |
| B | 145.88-232.95 | 182.75 | 1 |
| C | 114.58-187.29 | 145.83 | 1 |
| D | 87.14-141.71 | 110.73 | 1 |
| E | 2.35-4.42 | 3.27 | 1 |
| F | 128.09-204.91 | 160.86 | 1 |
| G | 1.43-2.81 | 2.04 | 1 |
| H | 117.92-195.06 | 151.36 | 0.5 |
| I | 124.77-200.69 | 157.02 | 0.49 |
| J | 88.78-146.1 | 113.73 | 1 |
| K | 66.37-115.22 | 88.32 | 1 |
| L | 66.31-109.73 | 85.29 | 1 |
| M | 3.13-8.92 | 5.71 | 1 |
| N | 101.87-168.21 | 130.49 | 1 |
| O | 81.02-132.17 | 102.87 | 1 |
| P | 45.84-84.94 | 63.25 | 1 |
| Q | 74.02-121.02 | 94.04 | 1 |
| R | 24.82-45.09 | 33.9 | 1 |
| S | 53.53-88.54 | 68.94 | 1 |
| T | 43.26-73.19 | 56.37 | 0.92 |
| U | 79.07-135.87 | 103.9 | 1 |
| V | 16.73-28.91 | 22.09 | 1 |
| X | 7.78-16.85 | 11.85 | 1 |
| Y | 9.82-17.37 | 13.15 | 1 |
| Z | 15.05-26.45 | 20.08 | 1 |

<sup>1</sup> Nodes present in Figure 3 and Figure 4.

**Supplementary file 1.** Test of substitution saturation of mitochondrial PCG.

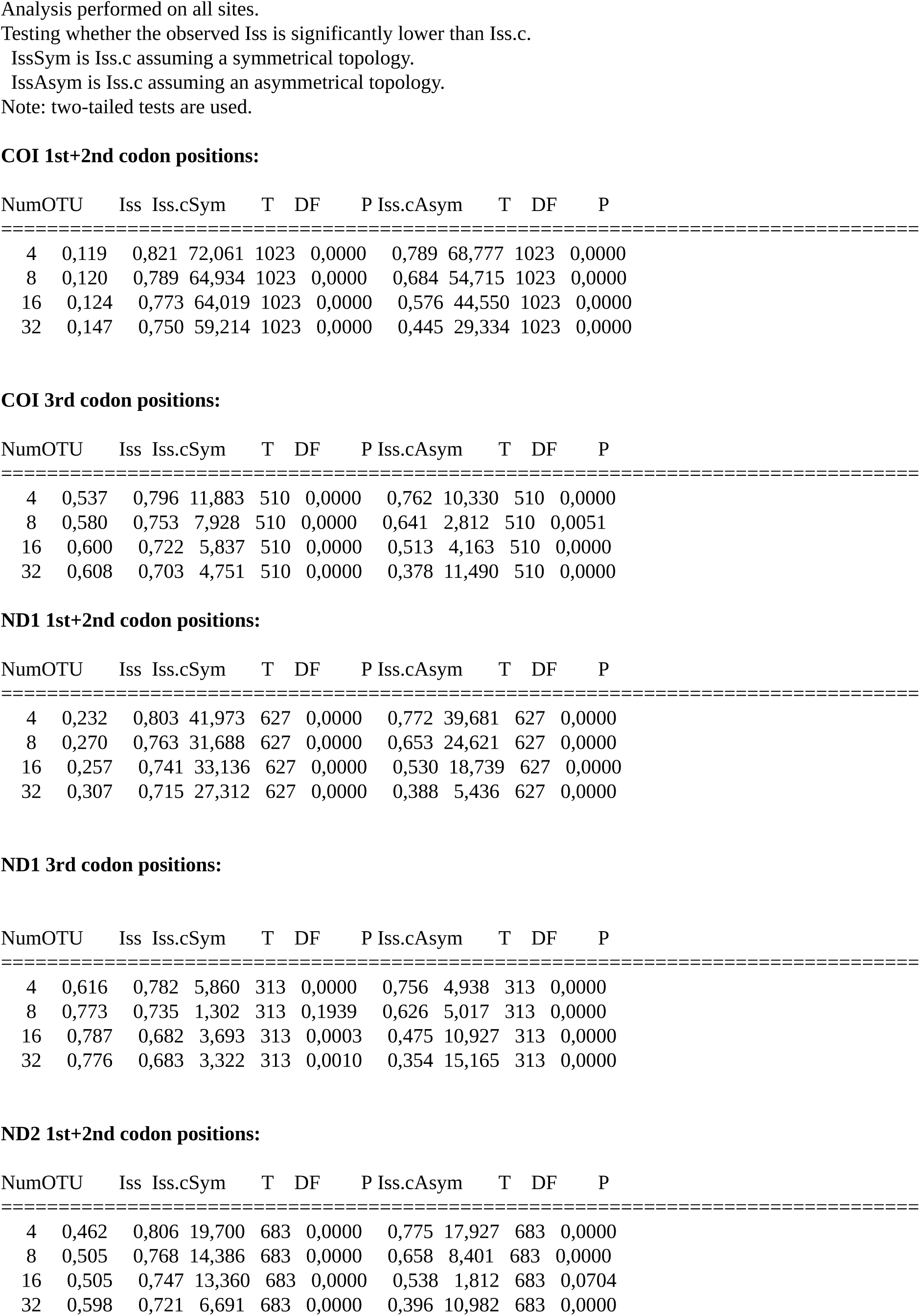
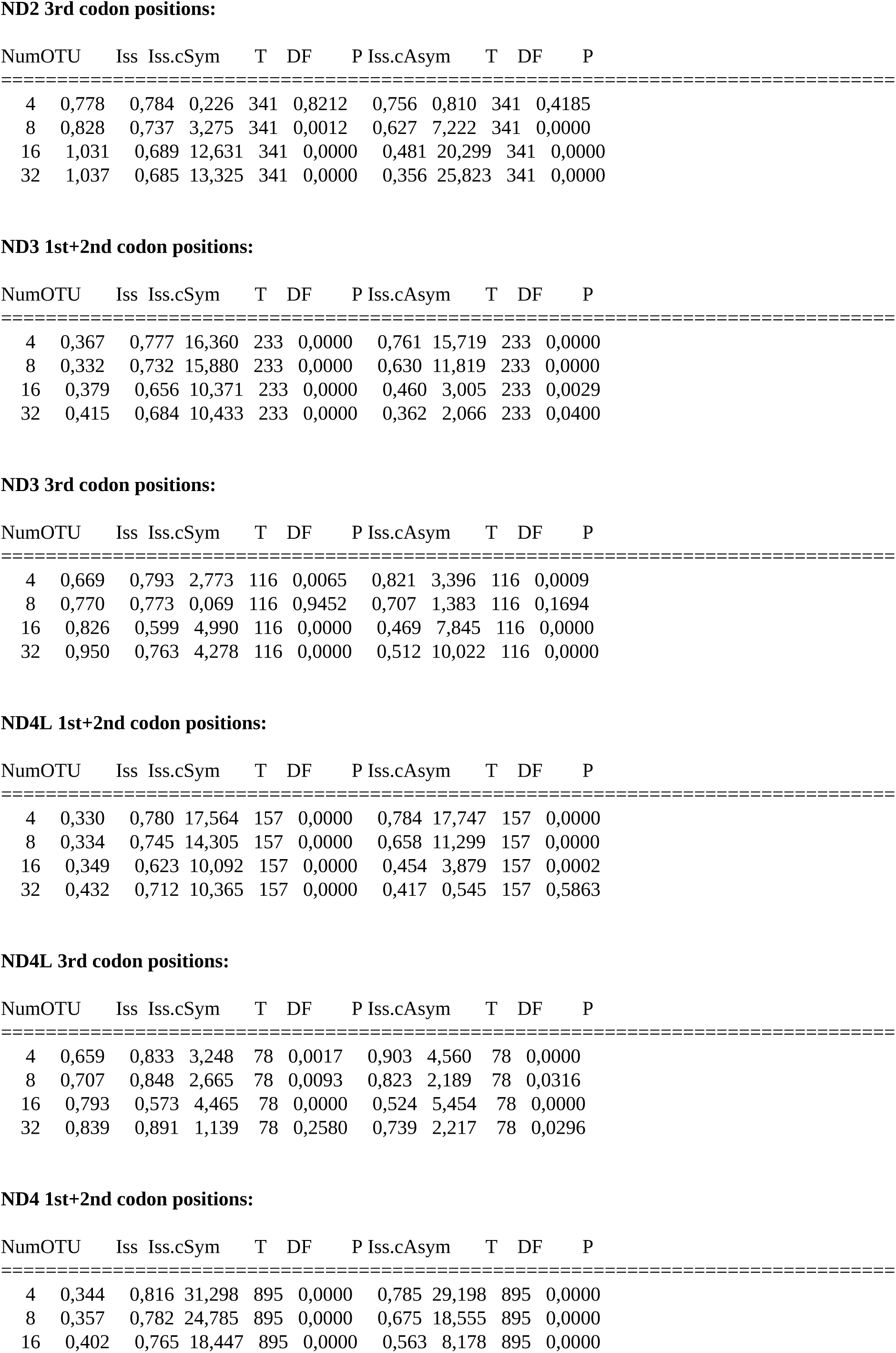
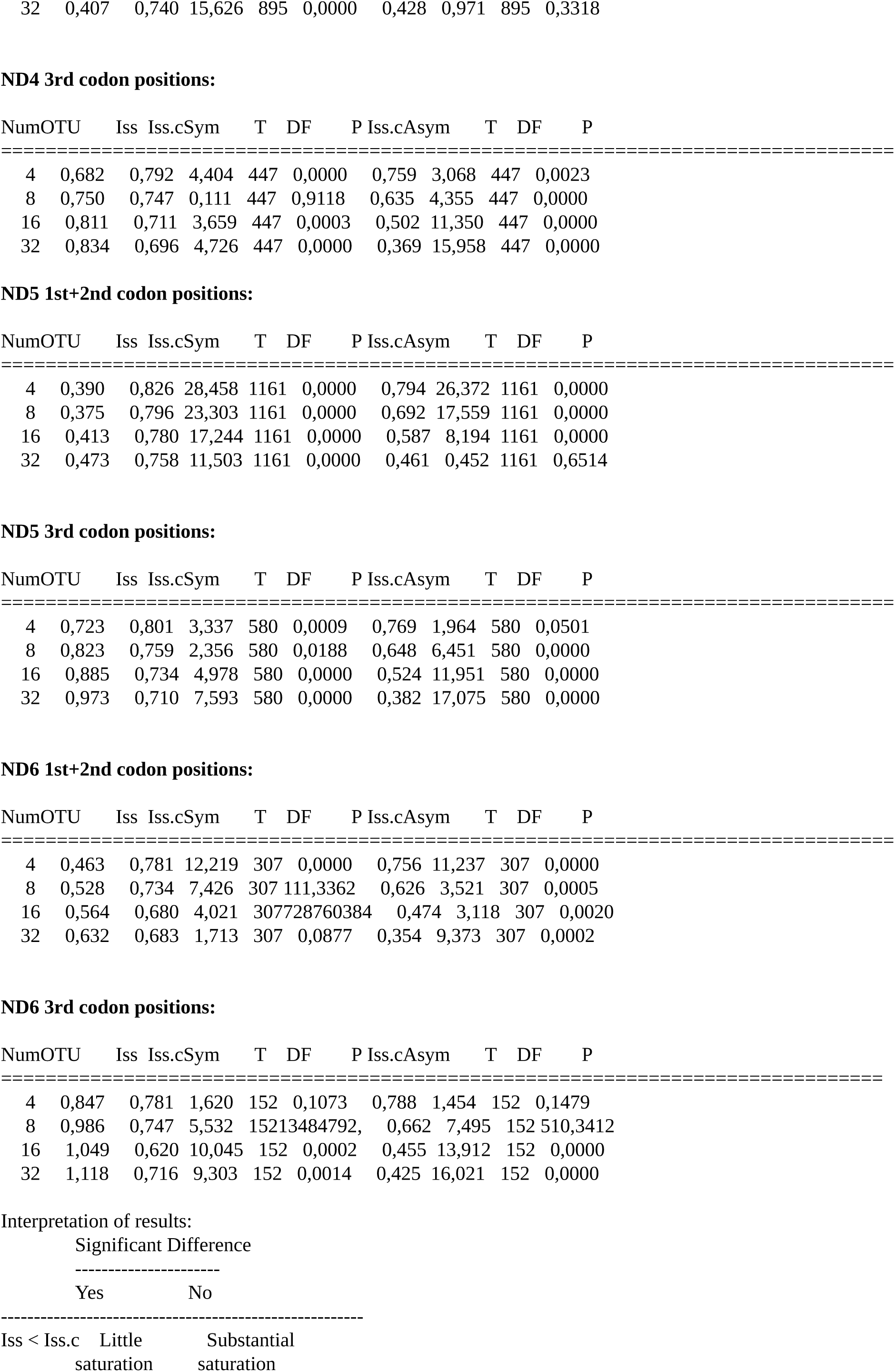

**Supplementary figure 1.**
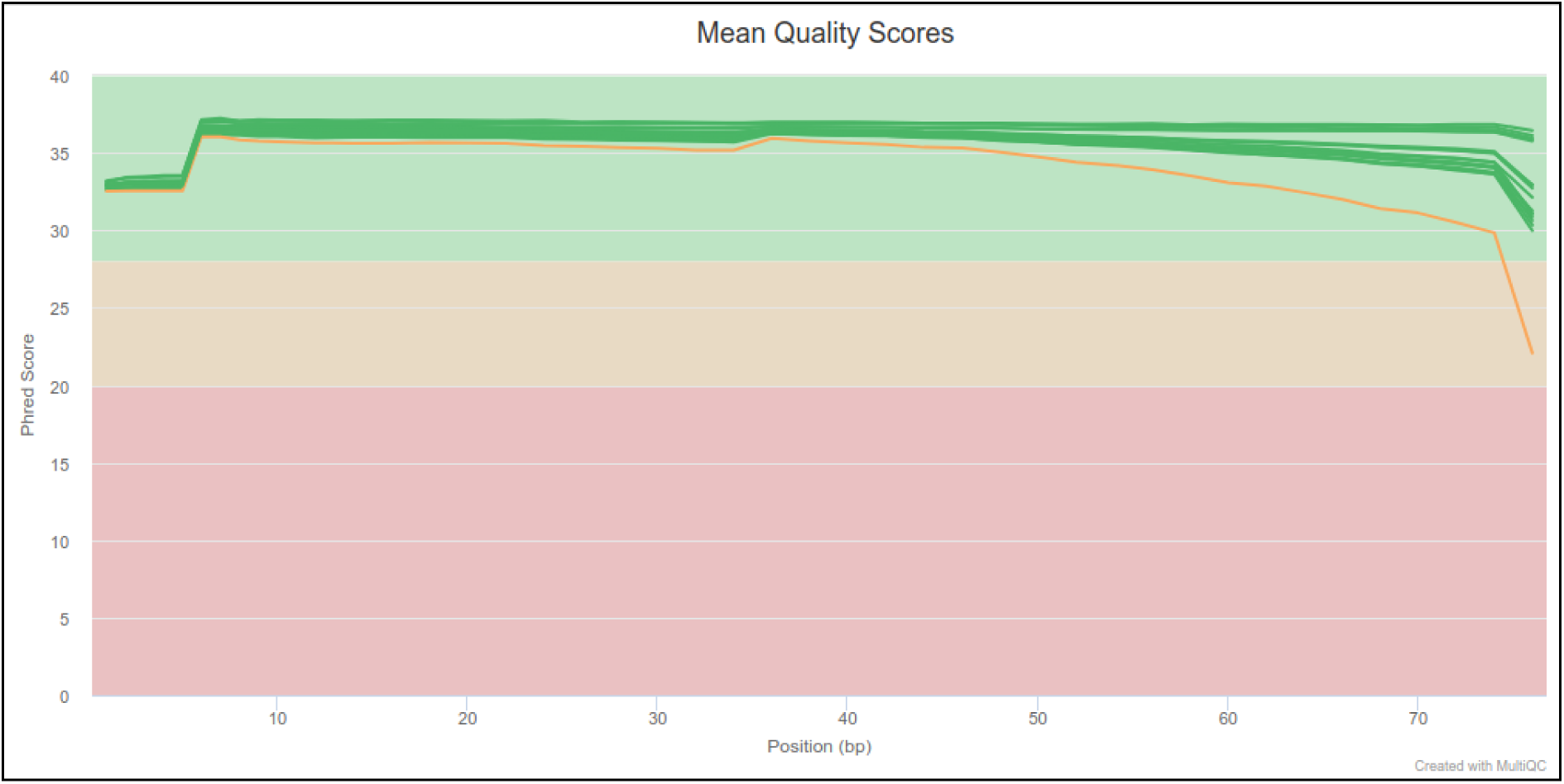
Quality of raw reads generated from mosquito samples. Figure generated from FasqQC results and summarized by MultiQC tool. The yellow line represents results from *Cq. albicosta* reads and green lines represent the remaining species.

**Supplementary figure 2.**
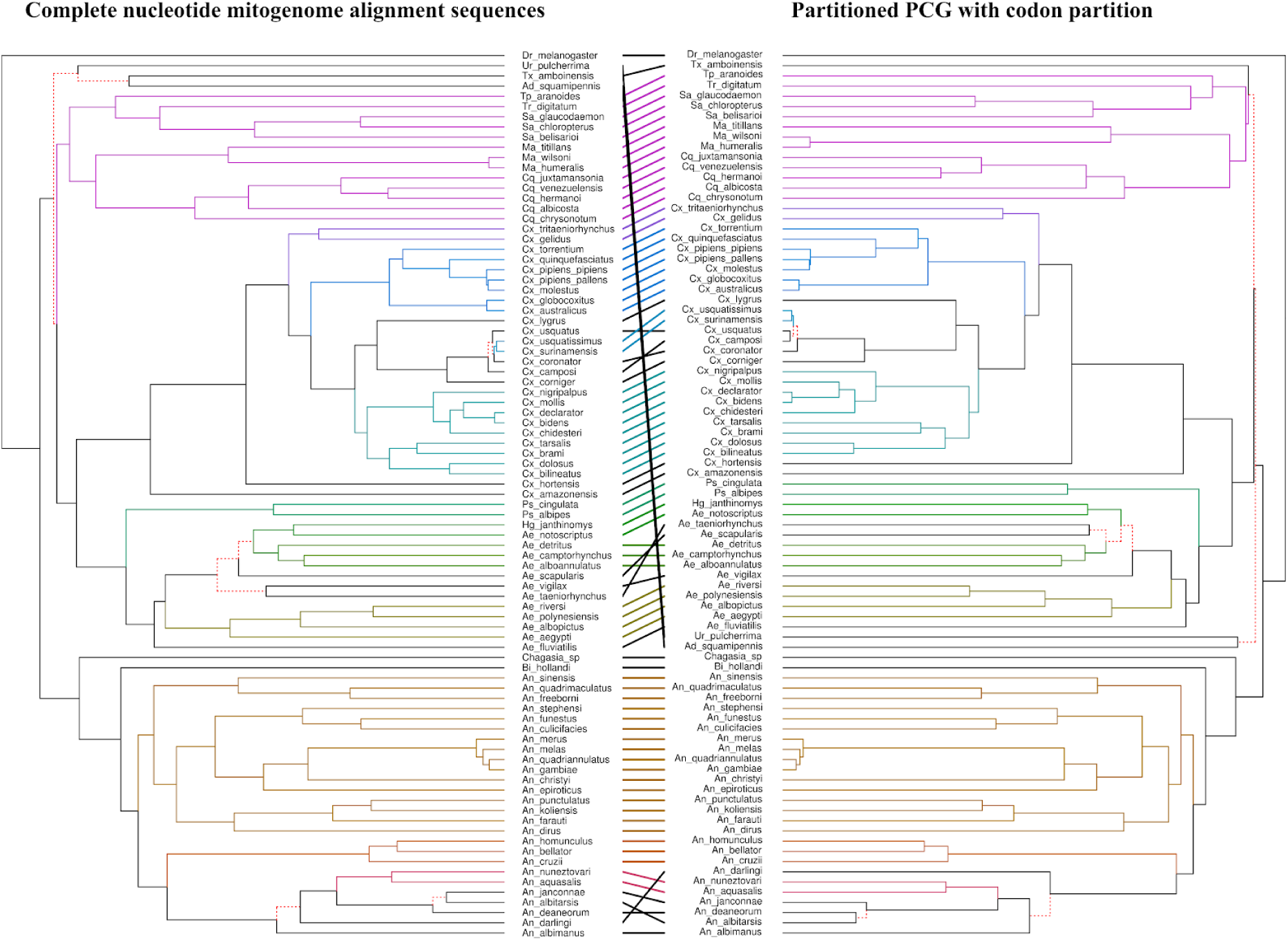
Comparative dendrogram topologies of trees obtained from different approaches using complete nucleotide mitogenomes and PCG with codon partition in the BEAST analysis. Dashed red lines show the branches that were different between the approaches used.

**Supplementary figure 3.**
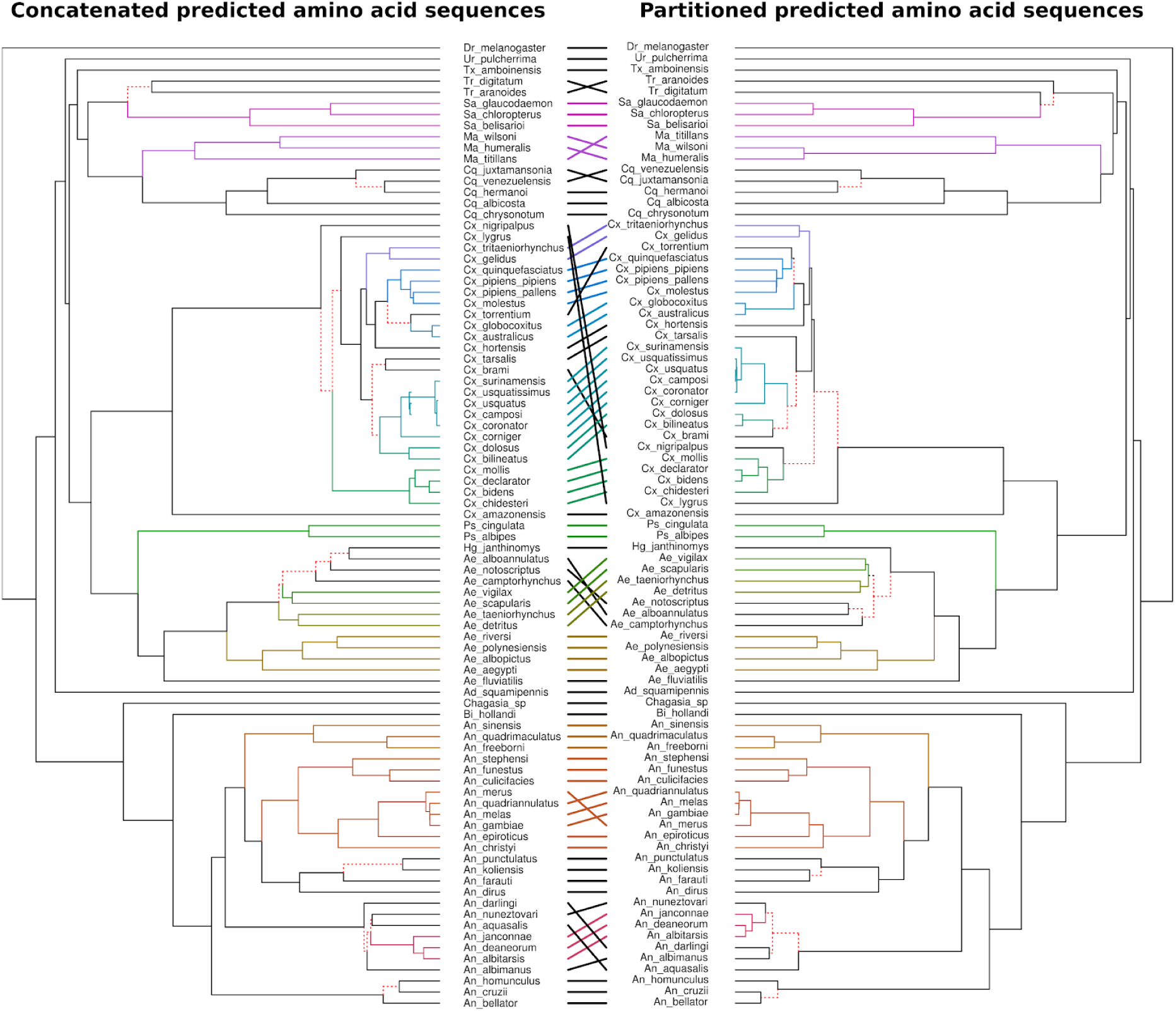
Comparative dendrogram topologies of trees obtained from different approaches using concatenated predicted amino acid sequences and partitioned predicted amino acid sequences in the BEAST analysis. Dashed red lines show the branches that were different between the approaches used.

**Supplementary figure 4.**
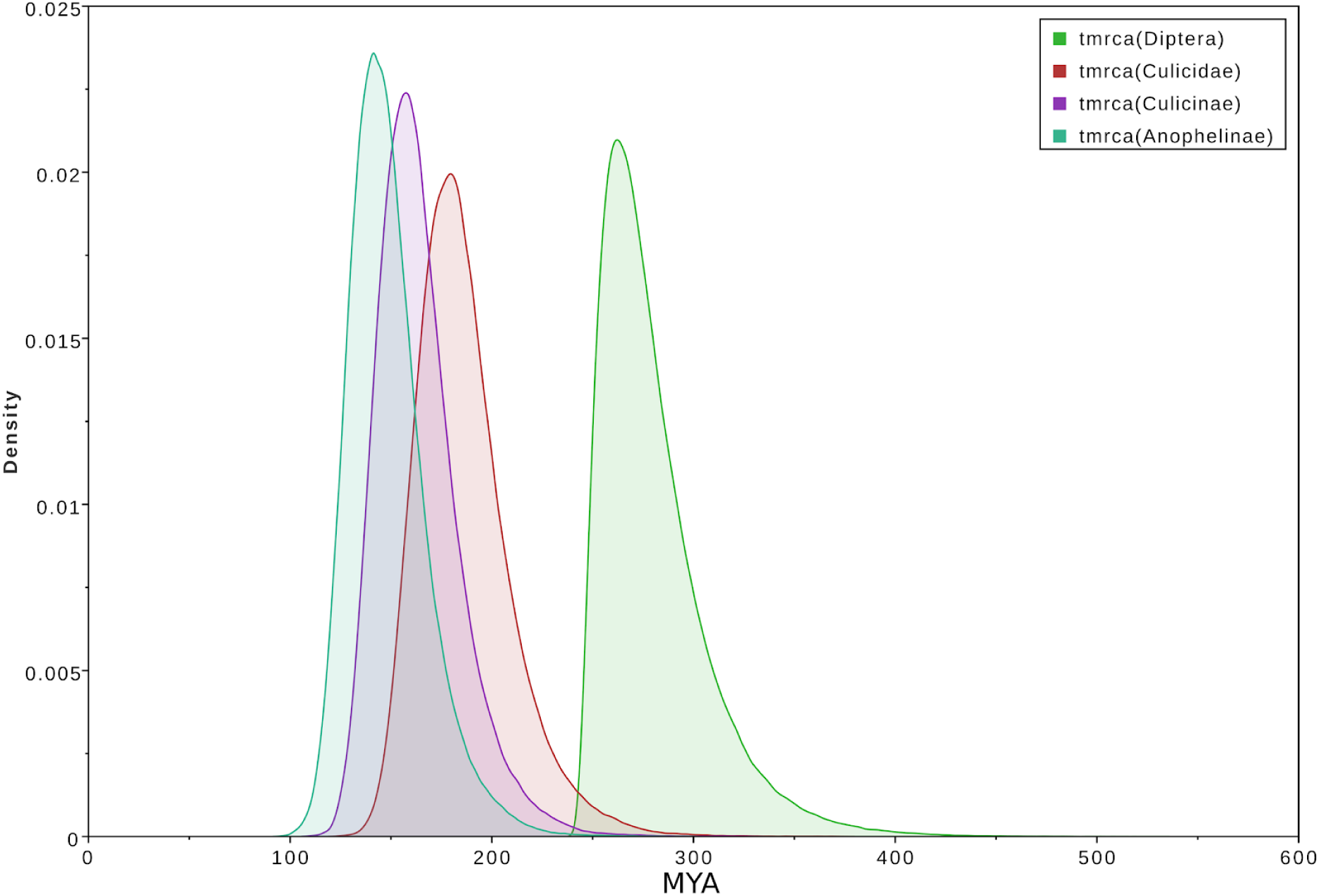
Density plot showing the 95% posterior distribution of ancestors time estimate based on the alignment of partitioned mitochondrial PCG with codon partition.

